# Neuroendocrine Control of Intestinal Regeneration Through the Vascular Niche in *Drosophila*

**DOI:** 10.1101/2024.09.10.612352

**Authors:** André B. Medina, Jessica Perochon, Cai Johnson, Sofia Polcowñuk, Yuanliangzi Tian, Yachuan Yu, Julia B. Cordero

## Abstract

Robust and controlled intestinal regeneration is essential for the preservation of organismal health and wellbeing and involves reciprocal interactions between the intestinal epithelium and its microenvironment. While knowledge of regulatory roles of the microenvironment on the intestine is vast, how distinct perturbations within the intestinal epithelium may influence tailored responses from the microenvironment, remains understudied. Here, we present previously unknown signaling between enteroendocrine cells (EE), vasculature-like trachea (TTCs), and neurons, which drives regional and global stem cell proliferation during adult intestinal regeneration in *Drosophila*.

Injury-induced ROS from midgut epithelial cells promotes the production and secretion of Dh31, the homolog of mammalian Calcitonin Gene-Related Peptide (CGRP), from anterior midgut EE cells. Dh31 from EE cells and neurons signal to Dh31 receptor within TTCs leading to cell autonomous production of the vascular endothelial growth factor (VEGF) and platelet-derived growth factor (PDGF)-like Pvf1. Tracheal derived Pvf1 induces remodeling of the tracheal stem cell niche and regenerative ISC proliferation through autocrine and paracrine Pvr/MAPK signalling, respectively. Interestingly, while EE Dh31 exerts broad control of ISC proliferation throughout the midgut, functions of the neuronal source of the ligand appear restricted to the posterior midgut. Altogether, our work has led to the discovery of a novel enteroendocrine/neuronal/vascular signaling network controlling global and domain specific ISC proliferation during adult intestinal regeneration.

## Introduction

The high regenerative nature of the intestinal epithelium, which is essential for organismal wellbeing, relies on tight coordination of interdependent stem cell intrinsic processes and multicellular paracrine signals (Hageman et al., 2020, Zhou and Boutros, 2023). Consistently, research on intestinal regeneration requires in vivo model systems that account for the complex inter-cellular and inter-tissue signaling between the gut epithelium and its natural microenvironment. Due its physiological sophistication, vast single cell molecular information and richness of tools for spatially and temporally restricted genetic manipulations, the fruit fly *Drosophila melanogaster* has been an excellent in vivo model system for the study of multi-organ communication including the gut (Medina et al., 2022).

Akin to the mammalian intestine, the adult *Drosophila* midgut regenerates through the action of stem cells (Micchelli and Perrimon, 2005, Ohlstein and Spradling, 2005) operating under the control of signals emerging from a rich microenvironment, collectively referred to as the intestinal stem cell (ISC) niche (Jiang et al., 2011, Jiang et al., 2009a, Amcheslavsky et al., 2014). In the mammalian gastrointestinal tract, diverse cellular types, including intestinal epithelium resident cells and cells from the associated mesenchyme provide physical support, contractility, and multiple stem cell niche factors such as Wnts, BMP, EGFs and R-Spondin, which instruct ISC self-renewal in homeostatic and challenging conditions (Kim et al., 2020, McCarthy et al., 2020, Stzepourginski et al., 2017, Niec et al., 2022, Palikuqi et al., 2022).

In addition to ISCs, the fly midgut epithelium consists of two subtypes of undifferentiated stem cell progeny, namely, enteroblasts (EBs) and pre-enteroendocrine cells, which are precursors of absorptive enterocytes (ECs) and secretory enteroendocrine cells (EEs), respectively (Chen et al., 2018). As in mammals, cells within the fly midgut epithelium and gut associated tissues, such as the visceral muscle (VM), terminal tracheal cells (TTCs) and enteric neurons (ENs) are components of the intestinal microenvironment that contribute to robust ISC proliferation and differentiation as part of a complete intestinal regeneration program (Medina et al., 2022, Petsakou et al., 2023).

A large body of research and cutting-edge technology has led to the discovery of the multiple components of the intestinal microenvironment and the myriad of stem cell niche factors they produce to instruct ISC function during tissue regeneration (Hageman et al., 2020, Kim et al., 2020, Harnack et al., 2019). However, less is known about how local and/or challenge-specific changes within the intestinal epithelium might influence the intestinal microenvironment to achieve robust and regenerative responses of the intestine, tailored to the environmental demands.

Secretory EE cells within the *Drosophila* and mouse intestine are a highly diverse, specialized and compartmentalized progeny of ISCs with the ability to sense a wide range of cues from within the intestine and the external environment (Guo et al., 2019, Gehart et al., 2019, Beumer et al., 2020, Lebrun et al., 2017). Signals received by EE cells are translated into the production of peptide hormones acting at short range, in an autocrine or paracrine fashion, or in a more classical endocrine or systemic mode, via their release into the circulation (Gribble and Reimann, 2016, Guo et al., 2022). The molecular and functional versatility of EE cells make them likely candidates to endow the gut with a highly refined regenerative response to the diversity of challenges faced by the intestinal epithelium. While nutrient regulated responses of EE cells and their constitutive function in the maintenance of intestinal homeostasis and growth have been recognized for decades (Scopelliti et al., 2014, Amcheslavsky et al., 2014, Martin et al., 2005, Drucker et al., 1996, Sasaki et al., 2001, Lin et al., 2022, Scopelliti et al., 2019), little is understood about the regulation and function of EE cells during intestinal regeneration. Recent datasets and publicly available toolkits for in situ gene expression analysis and independent genetic manipulation of EE peptide hormones and their cognate receptors, open a door for new and fundamental discoveries on the communication between EE cells and the intestinal microenvironment (Hung et al., 2020, Li et al., 2022, Liu et al., 2022b).

Here, we uncover multi-tissue signalling crosstalk between intestinal EE cells, gut associated vascular-like trachea and neurons, involving Calcitonin Gene-Related Peptide and PDGF/VEGF like factors and controlling global and domain specific intestinal regeneration in *Drosophila*.

## Results

### Intestinal damage promotes an evolutionary conserved increase in enteroendocrine cells

Damage-induced repair of highly regenerative tissues, such as the intestine, includes increased epithelial cell death and compensatory stem cell proliferation, resulting in significant changes in cellular dynamics in *Drosophila* (Amcheslavsky et al., 2009, Jiang et al., 2009b, He et al., 2018) and higher organisms (Parikh et al., 2019). Alterations in enteroendocrine (EE) cells are recognised features of intestinal damage (Dahly et al., 2003, Lebrun et al., 2017, Martin et al., 2005), pathogenic infection (Yang et al., 1990, Forbes et al., 2009, Bosi et al., 2005, Ovington, 1985, van Marle et al., 2013) and inflammatory bowel disease (IBD) (Worthington et al., 2018, Harrison et al., 2013). However, the aetiology and functional implications of EE cell changes to adult intestinal health and disease, remain largely unknown.

We analysed the proportion of EE cells in adult *Drosophila* midguts from mated females by staining with an antibody against the transcription factor, Prospero (Pros), and observed that damage to the midgut epithelium caused by oral exposure with the entomopathogen *Pseudomonas entomophila* (*Pe*), the DNA-damage agent bleomycin or the basal membrane disruptor dextran sodium sulphate (DSS), caused a significant increase in Pros positive EE cells (Figure 1A, 1B). Mouse intestinal samples from animals subject to DNA damage by whole body irradiation, or with inflammatory colitis induced by treatment with DSS also showed a noticeable increase in the number of Chromogranin A (ChgA) positive EE cells in the colon (Figure 1C, 1C’,1D). No significant change in EE cell numbers was observed in the small intestine (Figure 1C’’ and 1D). Taken together, our results suggest that increased EE cell proportion is a conserved feature of *Drosophila* and mouse intestines undergoing acute damage, which recapitulate observations made in the context of related human intestinal perturbations (Sciola et al., 2009, Zissimopoulos et al., 2014, Zhang et al., 2019). Next, we used *Drosophila* to investigate the nature of such EE phenotype and its functional significance to the intestinal response to damage.

**Figure 1:**
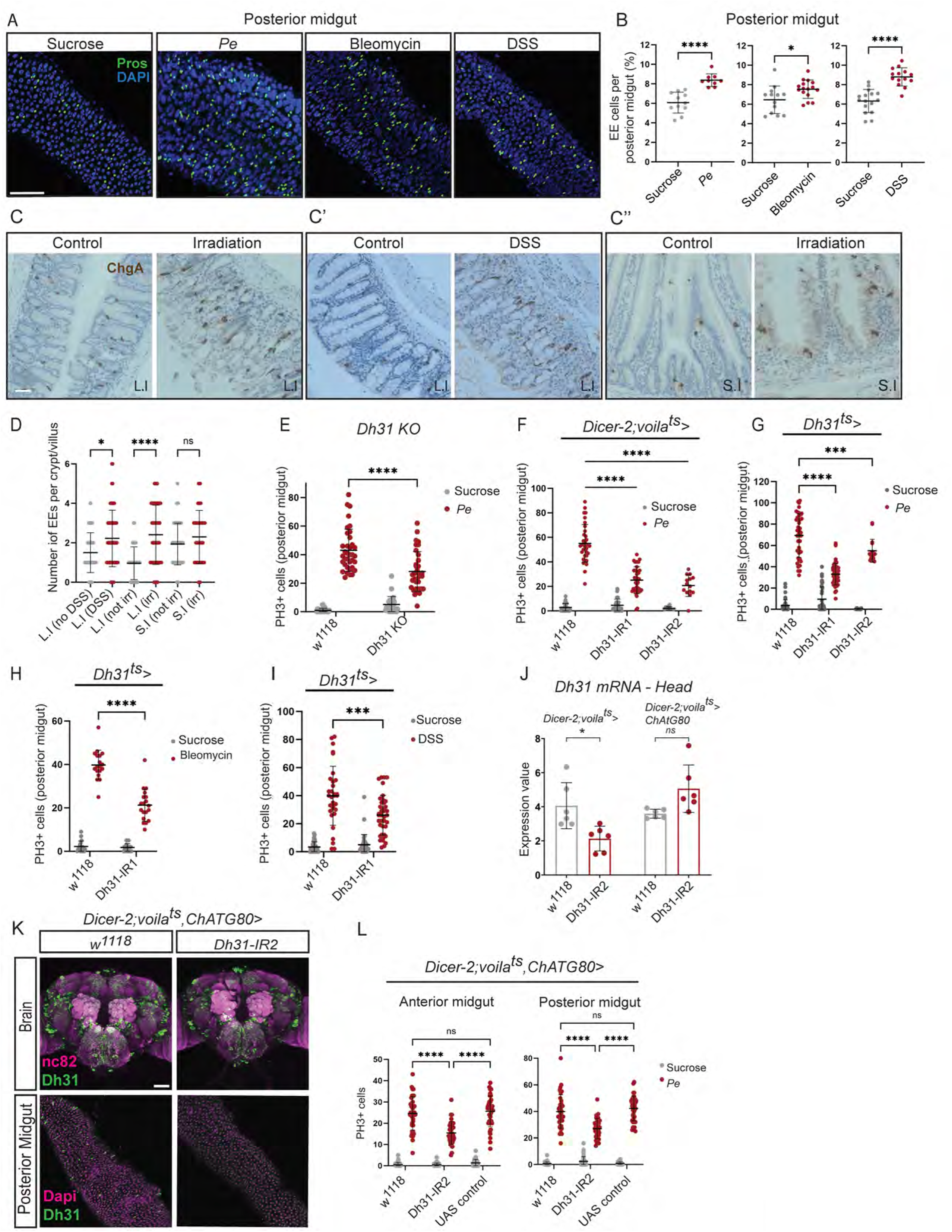
EE-cell derived peptide hormone Dh31 is required to fulfill the regenerative capacity of the midgut epithelium upon damage. (A) Immunofluorescence of EE cells labeled by Prospero (Pros; green) in the posterior midgut of flies fed with Sucrose, *Pe*, Bleomycin or DSS. (B) Quantification of EE cell proportion as in A. (C) Representative images of EE cells, labeled by Chromogranin A (ChgA; brown) immunohistochemistry, in the mouse large intestine (L.I) and small intestine (S.I). Control and whole-body irradiation (C) and control and DSS treatment (C’) in the L.I and control and whole-body irradiation in the S.I (C’’). (D) Quantification of EE cell proportion as in C, C’, C’’. (E) Quantification of PH3+ cells in Sucrose and *Pe* treated posterior midguts of Dh31 whole body knockout animals. (F) Quantification of PH3+ cells in Sucrose and Pe treated posterior midguts following Dh31 knockdown (*Dh31-*IR1 and *Dh31-*IR2) with *voil*á*-gal4*. (G, H, I) Quantification of PH3+ cells in posterior midguts treated with *Pe* (G), Bleomycin (H) and DSS (I) following *Dh31* knockdown (KD) (*Dh31*-IR1 and/or *Dh31*-IR2) in Dh31+ cells. (J) *Dh31* mRNA expression value in fly heads *Dh31* KD (*Dh31*-IR2) driven by *voil*á*-gal4* with or without *ChAT-gal80*. (K) Representative confocal images of Dh31 protein (green) levels in the brain and posterior midgut following *Dh31* KD (*Dh31*-IR2) with a gut specific driver (*voilá-gal4* with *ChAT-gal80)*. (L) Quantification of PH3+ cells in the anterior and posterior midgut of animals treated with Sucrose or *Pe* following Dh31 knockdown (*Dh31-*IR1 and *Dh31-*IR2) specifically in midgut EE cells (*voilá- gal4* with *ChAT-gal80)*. (B, D) Two-tailed unpaired Student’s t-test. (J) Ordinary one-way ANOVA. (E, F, G, H, I, L) Two-way ANOVA followed by Tukey’s multiple comparisons test. * (P<0.1) *** (P<0.001) **** (P<0.0001). Scale bar = 50 µm.

### Regulation of the EE-cell derived peptide hormone Dh31 is required to fulfill the regenerative capacity of the midgut epithelium upon damage

EE cells secrete peptide hormones that are constitutively required for the maintenance of ISC homeostasis and tissue growth in both fruit flies (Scopelliti et al., 2014, Amcheslavsky et al., 2014) and mammals (Drucker et al., 1996, Sasaki et al., 2001). However, the wide diversity and exquisite regionalization of EE cells in the gastrointestinal track (Beumer et al., 2020, Guo et al., 2019, Gehart et al., 2019) strongly suggest the potential for specialized and regulated responses of these cells to a wide range of stimuli. We therefore asked whether the global increase in EE cell numbers observed in damaged intestines may reflect on an EE cell sub-type specific response and/or function in tissue regeneration. We performed an RNA interference (RNAi) mini-screen to knockdown 9 different peptide hormones produced by EE cells using the temperature sensitive driver *voilá-Gal4^ts^*, which labels all EEs in the midgut (Scopelliti et al., 2014). We measured intestinal regeneration by counting the number of proliferative ISCs (PH3 positive) within the posterior midgut of mated females fed overnight with the pathogen *Pe*. We observed that knocking down the expression of *Dh31* caused the strongest and most consistent reduction in PH3 positive cells (Figure S1A, S1B), suggesting a preferential role for Dh31 producing EE cells in intestinal regeneration. Whole body *Dh31* loss of function and EE cell gene knockdown using independent RNAi lines (*Dh31-IR1* and *Dh31-IR2*) confirmed our initial observations that *Dh31* is required for intestinal regeneration (Figure 1E, 1F). Similarly, knocking down *Dh31* in *Dh31* expressing cells (*Dh31-Gal4^ts^*) in animals subject to intestinal damage by oral administration of *Pe*, bleomycin or DSS, resulted in significantly reduced regenerative ISC proliferation (Figure 1G-I).

Related to the neuroepithelial characteristics of EE cells, peptide hormones secreted by those gut cells are also produced by neuronal cells in the brain. Therefore, genetic drivers broadly targeting EE cells, including *voilà-Gal4* and *Dh31- Gal4* are expressed in both, the midgut and neurons (Scopelliti et al., 2014, Lin et al., 2022). To address EE specific functions of Dh31, we screened for a combination of genetic tools that would reduce *Gal4* expression and therefore impair gene targeting within neurons, without compromising gene manipulations in EE cells (Figure S1C). We found that the cholinergic line, *ChAT-Gal80* (Kitamoto, 2002) strongly decreased GFP expression driven by *voilá-Gal4* in central and peripheric neurons, with no effect on the midgut (*Dicer-2;voilá^ts^;Chat-gal80>GFP*; Figure S1C). Using such genetic combination we observed that, while a significant reduction of *Dh31* expression was observed in the heads of *voilá^ts^* driven *Dh31-*RNAi animals, *Dh31* knockdown using *voilá^ts^,ChAT-Gal80* did not affect brain gene expression levels (Figure 1J). Consistently, we observed no reduction in Dh31 immunostaining in either the brain or ventral nerve cord (VNC) of *voilà^ts^,ChAT-Gal80*>*Dh31-IR* animals, while midgut protein signal was undetectable (Figure 1K, S1D). Importantly, gut specific *Dh31* knockdown was sufficient to impair regeneration in damaged midguts as revealed by reduced ISC proliferation in the anterior and posterior midgut (Figure 1L). However, overexpression of *Dh31* or activation of Dh31^+^ EE cells through expression of the mammalian capsaicin receptor (VR1) was not sufficient to promote ISC proliferation in the absence of damage (Figure S1E, S1F). We noted that, in addition to EE cells, Dh31 was expressed in a subset of enteric neurons, most predominantly, in hindgut innervating neurons (Cognigni et al., 2011) (Figure 2A). Pan-neuronal knockdown of *Dh31* under the control of *nSyb-gal4* (*nSyb^ts^*) decreased Dh31 levels in hindgut enteric neurons (Figure 2A’) and gene expression in the head (Figure 2B) but did not affect Dh31 immunostaining in EE cells (Figure 2A’). Interestingly, neuronal *Dh31* knockdown resulted in reduced regenerative ISC proliferation restricted to the posterior midgut (Figure 2C). Altogether, these results indicate that EE cell derived Dh31 is broadly needed for damage induced regeneration of the adult *Drosophila* midgut. Furthermore, an additional source of the neuropeptide, produced by neurons, contributes locally to posterior midgut regeneration.

**Figure 2:**
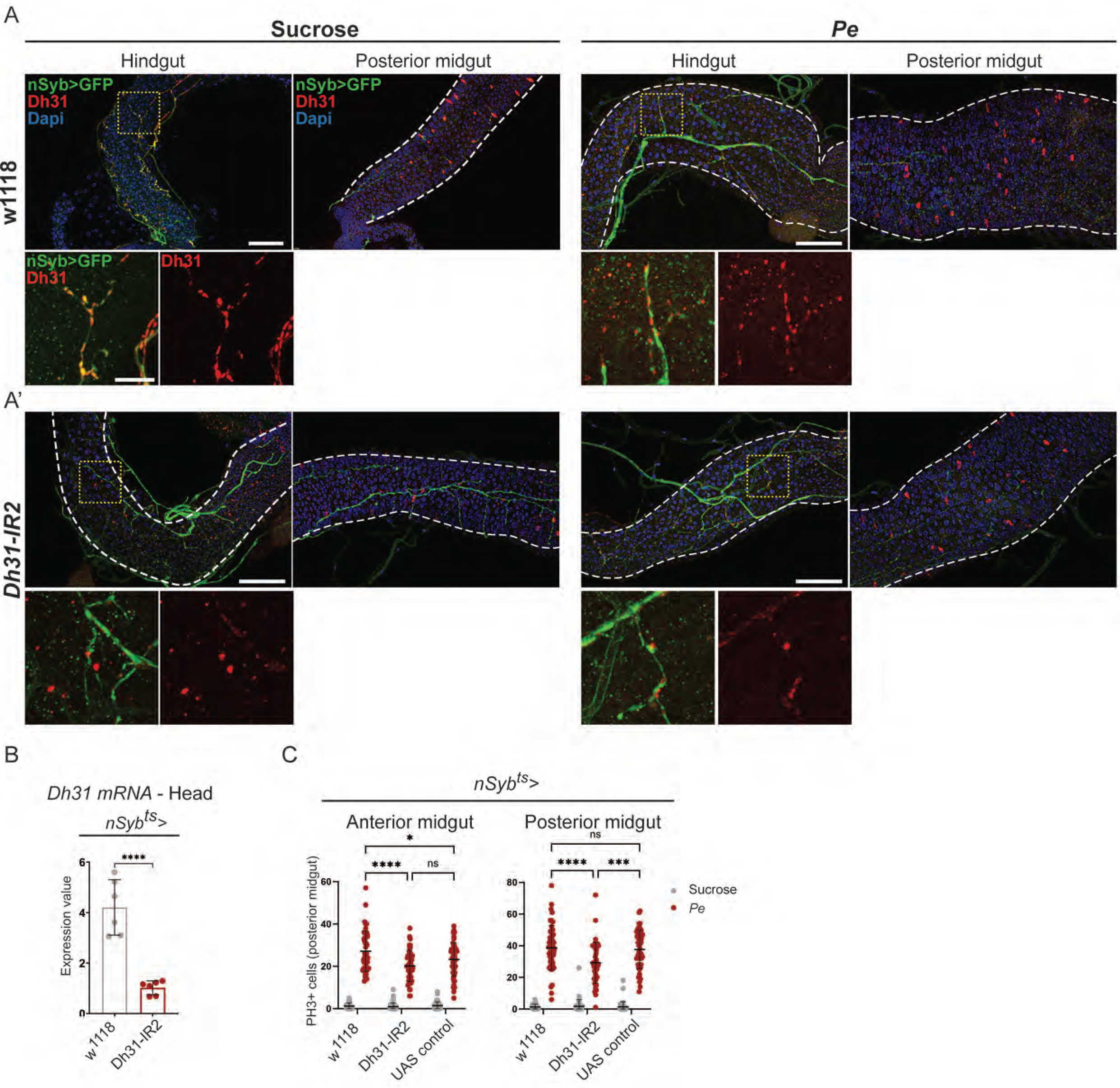
Neuronal-derived peptide hormone Dh31 is required to fulfill the regenerative capacity of the posterior midgut epithelium upon damage. (A, A’) Confocal images of *nSyb>GFP* expression (green) in the adult *Drosophila* hindgut and posterior midgut co-stained with and anti-Dh31 (red). Dashed squares indicate the magnified view presented below each full-size figure panel. Images corresponds to hindguts and midguts from control animals *w^1118^*(A) and animals with *Dh31* KD *(Dh31*-IR2) under the neuronal driver *nSyb-gal4^ts^* (A’) and fed with Sucrose (A, A’; left panels) or *Pe* (A, A’; right panels). (B) RT-qPCR of head *Dh31* mRNA levels in control animals or animals with *Dh31* KD *(Dh31*-IR2) under the neuronal driver *nSyb-gal4^ts^*. (C) Quantification of PH3+ cells in sucrose and *Pe* treated anterior and posterior midguts following *Dh31* knockdown (*Dh31*-IR1) with *nSyb- gal4*. Gray dots represent individual cells and red dots represent the average value per midgut. (B) Two-tailed unpaired Student’s t-test. (C) Two-way ANOVA followed by Tukey’s multiple comparisons test * (P<0,04), ** (P<0,008), *** (P<0.001), **** (P<0.0001). Scale bar = 50 µm.

### Dh31 production is regulated by intestinal damage

We next assessed the distribution of Dh31^+^ EE cells in the midgut using a *Dh31-Gal4>mCD8-GFP* gene expression reporter and an antiserum against Dh31 protein. Consistent with previous reports (Veenstra et al., 2008, Benguettat et al., 2018), we detected high levels of Dh31 immune-labelling and gene expression in posterior midgut regions R4c and R5 (Figure 3A, B’, C’, D and S2A), and a low number of cells with weaker protein and gene expression signal in the rest of the homeostatic midgut (Figure 3A-D and S2A). On the other hand, oral infectious challenge to the intestinal epithelium with *Pe*, resulted in strong increase in immuno- labelling of Dh31^+^ EE cells and gene expression in the anterior midgut (Figure 3B-D), with no significant changes detected in the posterior midgut (Figure 3B’,C’,D). Altogether, these results suggest domain specific upregulation of *Dh31* mRNA and protein in response to pathogenic damage, which is restricted to the anterior midgut.

**Figure 3:**
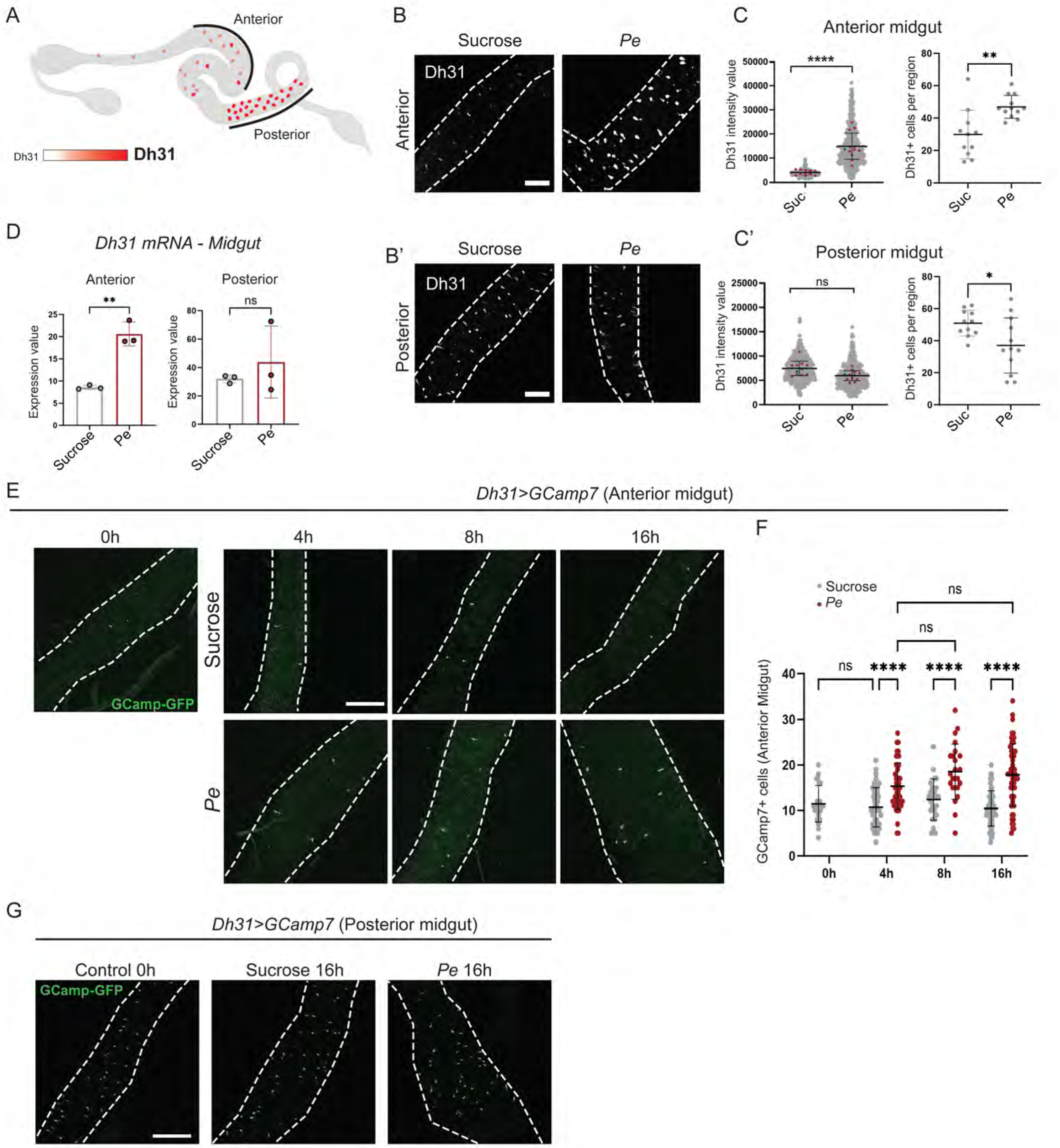
Dh31 production is regulated by intestinal damage. (A) Schematic view of Dh31 expression pattern in the midgut. Scale from white (less expression) to red (more expression). (B) Expression levels of Dh31 detected by immunostaining upon treatment with Sucrose or *Pe* in the anterior (B) and posterior midgut (B’). (C, C’) Quantifications of Dh31 signal intensity levels and Dh31+ cell numbers in the anterior (C) and posterior midgut (C’). (D) *Dh31* mRNA expression value in anterior and posterior midguts from Sucrose and *Pe* treated animals. (E) Ex-vivo visualization of GCamp7c (green) expression in Dh31+ cells in the anterior midgut of flies treated with Sucrose or *Pe* for 0h, 4h, 8h and 16h. (F) Quantification of GCamp7 + cells as in E. (G) Ex-vivo visualization of GCamp7 expression in Dh31+ cells in the posterior midgut of flies untreated or treated with Sucrose or *Pe* for 16h. Dashed line delineate the edges of the midgut. (C, C’, D) Two-tailed unpaired Student’s t-test. (F) Two-way ANOVA followed by Tukey’s multiple comparisons test. * (P<0.1) ** (P<0.002) **** (P<0.0001). Scale bar = 100 µm.

EE cells store peptide hormones into vesicles and secrete them locally or systemically into the haemolymph following activation by specific stimuli in a process that involves increase in intracellular Calcium (Ca^2+^) (Scopelliti et al., 2018, Lin et al., 2022). To further interrogate the nature of upregulated Dh31 upon midgut damage (Figure 3A-D), as to whether this may reflect increased peptide retention or higher peptide production and secretion, we measured Dh31^+^ EE cell activity *ex vivo* by assessing cytoplasmic Ca^2+^ levels using GCaMP7, a genetically encoded calcium sensor. We expressed GCaMP7 in Dh31^+^ EE cells after different periods of 5% sucrose or *Pe* feeding. We detected increased Dh31^+^ cell activity in the anterior midgut as soon as 4h of infection, which remained consistently high after 8h and 16h of infection (Figure 3E, F). Co-expression of GCaMP7 and the nuclear reporter RedStinger, using a *Dh31* knock-in *Gal4* line *Dh31-T2A-Gal4*, showed that almost all Dh31^+^ EE cells were activated in the anterior midgut after 16h of *Pe* infection when compared to sucrose fed animals (Figure S2B). We did not detect changes in basal levels of Ca^2+^ in Dh31^+^ EE cells within the posterior midgut, which remained constitutively high regardless of treatment (Figure 3G). Taken together, our results show a compartmentalized regulated response of EE cells to intestinal damage, resulting in increased production and secretion of Dh31 from the anterior compartment of the adult *Drosophila* midgut.

### Gut epithelial derived reactive oxygen species induce Dh31 production in *Drosophila*

Intestinal damage triggered by pathogenic bacteria infection generates high levels of reactive oxygen species (ROS) in the gut lumen, mainly via enterocytes (ECs), as a protective mechanism from the host against the pathogen (Morris and Jasper, 2021). Dh31^+^ EE cells are responsive to extracellular ROS generated by short-term bacterial infection (Benguettat et al., 2018). Blocking ROS in the adult midgut by feeding *Pe* infected animals with the antioxidant *N-acetyl cysteine* (NAC) or by genetically inhibiting ROS production in ECs via DUOX knockdown led to a significant reduction in EE cell Dh31 immunolabelling in the anterior midgut (Figure 4A-D). Furthermore, feeding animals with a sucrose solution containing H_2_O_2_, a form of ROS commonly produced during bacterial infection, increased EE cell Dh31 immunolabelling specifically within the anterior midgut (Figure 4E-H). Therefore, ROS produced by midgut epithelial cells is necessary and sufficient to induce Dh31 production in EE cells of the adult anterior midgut.

**Figure 4:**
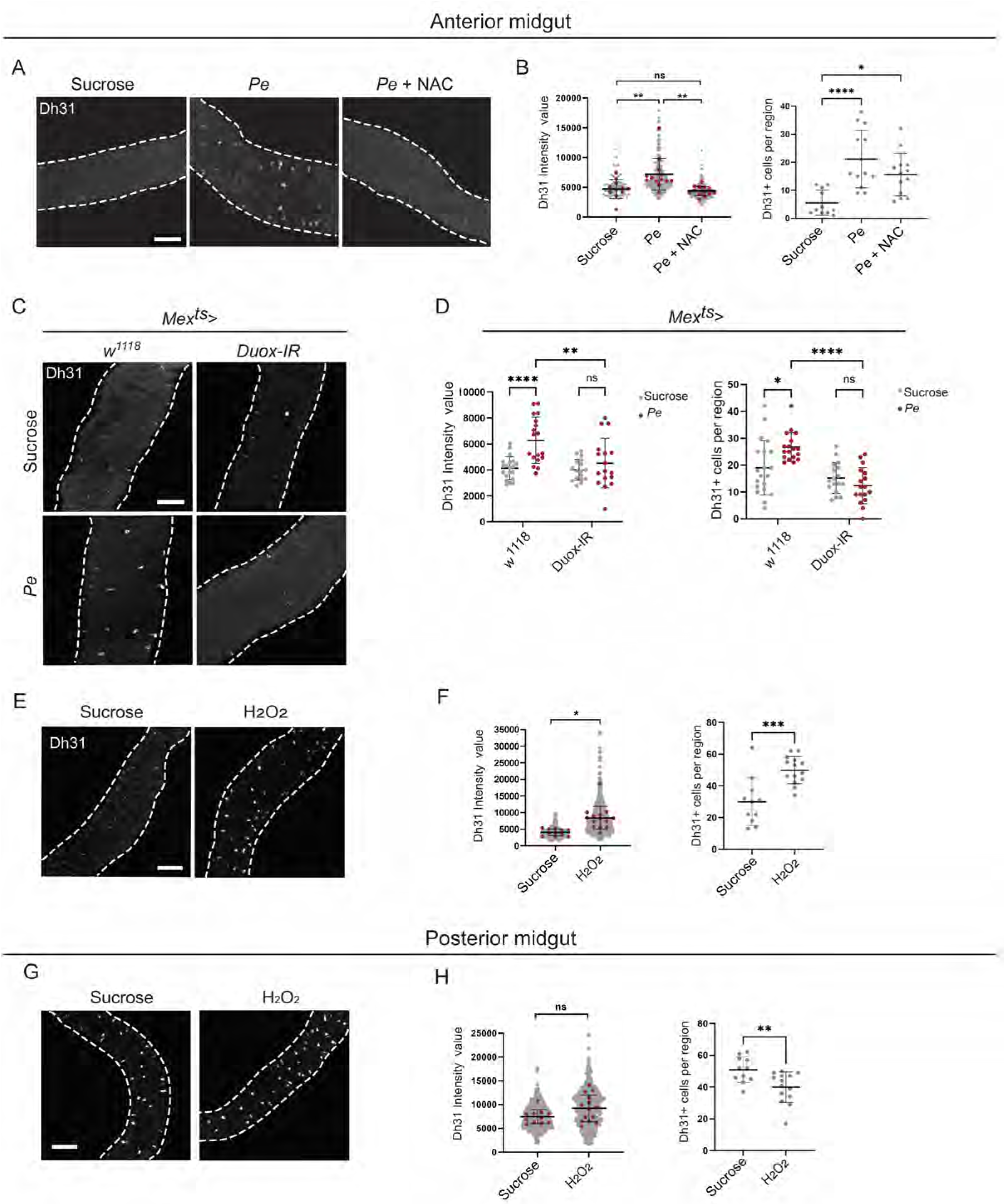
**Gut epithelial derived reactive oxygen species induce Dh31 production in *Drosophila***. (A) Expression levels of Dh31 (grey) detected by immunostaining upon treatment with Sucrose, *Pe* or *Pe* + NAC in the anterior midgut. (B) Quantifications of Dh31 signal intensity value and Dh31+ cell numbers in anterior midguts as in A. (C) *Duox* knockdown in ECs (*Mex-gal4*) and effect on Dh31 protein levels (grey) in anterior midguts of flies treated with Sucrose and Pe. (D) Quantifications of Dh31 signal intensity value and Dh31+ cell numbers in anterior midguts as in C. (E, G) Expression levels of Dh31 (grey) upon treatment with Sucrose or hydrogen peroxide (H_2_O_2_) in anterior (E) and posterior (G) midguts. (F, H) Quantifications of Dh31 signal intensity value and Dh31+ cell numbers in anterior midguts as in E and G, respectively. Dashed lines delineate the edges of the midgut. (B) Ordinary one-way ANOVA and (D) Two-way ANOVA followed by Tukey’s multiple comparisons test. (F) Two-tailed unpaired Student’s t-test. * (P<0.04), ** (P<0.008), *** (P<0.003), **** (P<0.0001). Scale bar = 100 µm.

### Tracheal Dh31 receptor signaling is required for ISC proliferation during midgut regeneration

To understand how damage-induced Dh31 signals to promote intestinal regeneration, we next investigated the expression of the Dh31 receptor (Dh31R). Single cell and bulk RNAseq as well as loss of functional experiments, have previously demonstrated the presence of *Dh31R* in midgut ECs, EEs and the visceral muscle (VM) (Dutta et al., 2015, Benguettat et al., 2018, Hung et al., 2020). We used a *Dh31-R* knock-in *Gal4* reporter, which recapitulates the endogenous expression pattern of the *Dh31-R* gene (Deng et al., 2019), to express GFP (*Dh31- R>GFP)* and therefore visualize receptor expression in situ. Consistent with previous reports (Dutta et al., 2015, Benguettat et al., 2018, Hung et al., 2020), we observed GFP expression in VM and ECs in the anterior midgut (Figure 5A and S3A), we also observed expression in uncharacterised enteric neurons of the crop and anterior midgut, and PDF expressing enteric neurons (Cognigni et al., 2011) in the posterior midgut (Figure 5A and S3A). Interestingly, we also observed *Dh31-R>GFP* expression in terminal tracheal cells (TTCs) though out the full length of the midgut (Figure 5A, B). We used inducible RNAi lines to knockdown *Dh31R* from various midgut compartments and midgut associated tissues, including PDF producing neurons (*Pdf^ts^>Dh31R-IR1),* visceral muscle (*how^ts^>Dh31R-IR1)*, ECs (*Mex^ts^>Dh31R-IR1)* and in TTCs (*dSRF^ts^>Dh31R-IR1)* (Figure S3B, C). While mild reduction in intestinal regeneration was observed upon *Dh31R* knockdown in the VM and ECs (Figure S3D,E), receptor downregulation in TTCs resulted in the most robust and consistent impairment of regenerative ISC proliferation throughout the midgut of *Pe* infected animals (Figure 5C, D and S3F) or fed with bleomycin (Figure 5E). Therefore, we focused the rest of our work on studying how Dh31 signalling to its tracheal receptor regulates intestinal regeneration.

**Figure 5:**
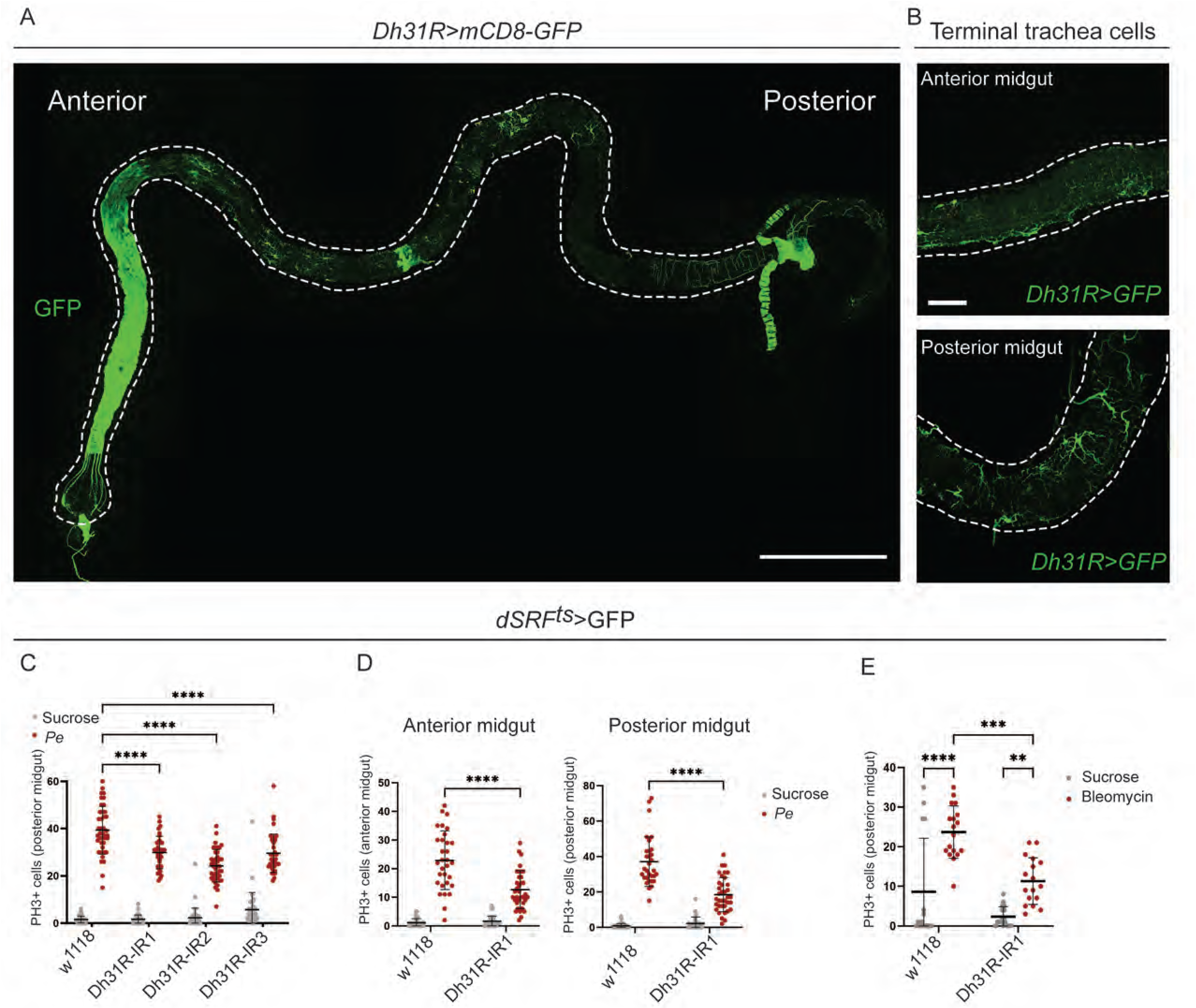
Tracheal Dh31 receptor (Dh31R) signaling is required for ISC proliferation during midgut regeneration. **(A)** Expression pattern of *Dh31R* knock- in *gal4* visualized by GFP (green) in the whole midgut. Scale bar = 1000 µm . (B) Closer view of additional anterior and posterior midgut regions highlighting *Dh31R>GFP* expression in TTCs. Scale bar = 100 µm (C) Quantification of PH3+ cells in the posterior midgut of animals treated with Sucrose or *Pe* following *Dh31R* knockdown (*Dh31R-*IR1, *Dh31R-*IR2 and *Dh31R-*IR3) in TTCs. (D) Quantification of PH3+ cells in the anterior and posterior midgut of animals treated with Sucrose or *Pe* following Dh31R knockdown (*Dh31R-*IR1) in TTCs. (E) Quantification of PH3+ cells in the posterior midgut of animals treated with Sucrose or Bleomycin following *Dh31R* knockdown (*Dh31R-*IR1) in TTCs. (C-E) Two-way ANOVA followed by Tukey’s multiple comparisons test. ** (P<0.009), *** (P<0.002), **** (P<0.0001). TTCs; terminal tracheal cells.

### Dh31R signaling affects TTCs remodeling and MAPK activation in the regenerating midgut

Adult terminal tracheal cell plasticity and the production of stem cell niche factors by the trachea are essential to support the regenerative function of ISCs following midgut injury (Perochon et al., 2021, Tamamouna et al., 2021). Downregulating *Dh31R* in TTCs (*dSRF^ts^>Dh31R-IR*) led to significantly diminished TTC remodeling in *Pe* infected midguts (Figure 6A, B). These results suggest that the activation of Dh31/Dh31R signaling within adult gut trachea and its contribution to midgut regeneration, involves the induction of TTC remodeling. ROS inducible production of the *Drosophila* FGF-like ligand Branchless (Bnl) by the trachea and intestinal epithelium, drives regenerative ISC proliferation through paracrine activation of the FGF receptor Breathless (Btl) and downstream MAPK signaling in ISCs (Perochon et al., 2021, Tamamouna et al., 2021). While we observed significant downregulation of MAPK activation in ISCs from injured midguts with impaired tracheal Dh31R signaling (*dSRF^ts^>Dh31R-IR)* (Figure 6C, D), there was no impact on *bnl* upregulation (Figure S4A,B). These data suggest that Dh31R dependent tracheal remodeling and MAPK activation within ISCs is unlikely to be mediated by Bnl/Btl signaling.

**Figure 6:**
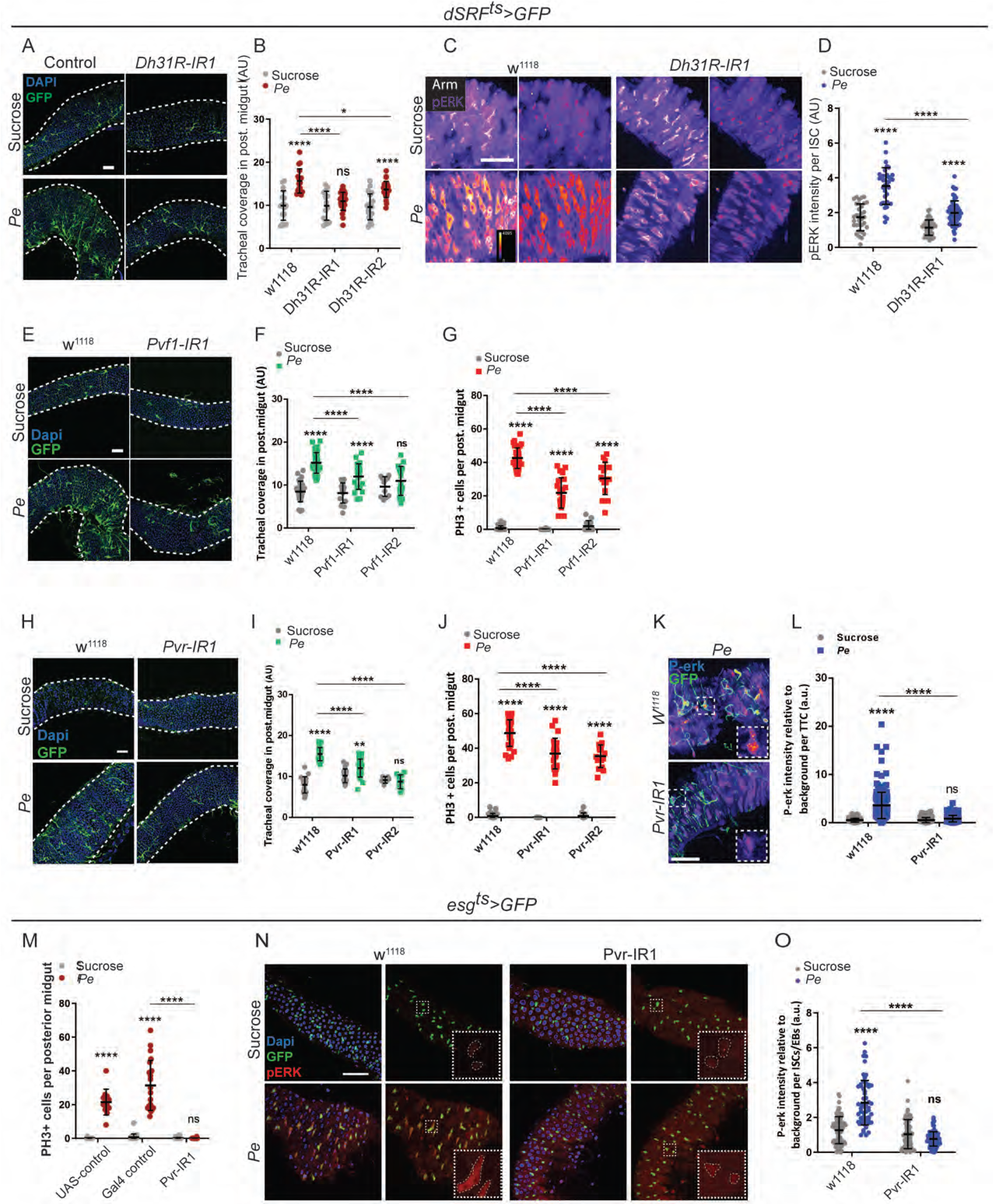
Dh31R signaling affects TTCs remodeling and MAPK activation in the regenerating midgut through activation of Pvf1/Pvr signaling. (A) Confocal imaging of tracheal coverage (green) in posterior midguts following *Dh31R* knockdown (*Dh31R*-IR1) in TTCs in animals treated with Sucrose or *Pe*. (B) Quantification of tracheal coverage in midguts as in A. (C) Representative confocal images of p-ERK (red) immunostaining in ISCs/EBs (small cells stained with anti-Armadillo (Arm; white) from posterior midguts upon *Dh31R* KD (*Dh31R*-IR1) in TTCs of animals treated with Sucrose or *Pe*. p-ERK is indicated in a fire scale (blue = low, yellow = high) to facilitate signal intensity visualization. (D) Quantification of p-ERK signal intensity in ISCs/EBs of posterior midguts as in C. (E) Confocal imaging of tracheal coverage (green) in posterior midguts following *Pvf1* knockdown (*Pvf1-*IR1) in TTCs and treated with Sucrose or *Pe*. (F, G) Quantification of tracheal coverage (F) and PH3+ ISCs (G) in posterior midguts a in E. (H) Confocal imaging of tracheal coverage (green) in posterior midguts following *Pvr* knockdown (*Pvr-*IR1) in TTCs and treated with Sucrose or *Pe*. (I, J) Quantification of tracheal coverage (I) and PH3+ ISCs (J) in posterior midgut a in H. (K) Representative confocal images of p- ERK immunostaining (red) in the posterior midgut upon *Pvr KD* (*Pvr*-IR1) in TTCs (green) after treatment with *Pe.* (L) Quantification of p-ERK signal in TTCs of midguts as in K after Sucrose or *Pe* treatment. (M) Quantification of PH3+ ISCs in posterior midguts upon *Pvr KD* (*Pvr*-IR1) in ISC/EBs. (N) Representative confocal images of p-ERK (red) immunostaining in ISCs/EBs (green) from posterior midguts of animals treated with Sucrose or Pe following *Pvr* knockdown (*Pvr-*IR1) in ISCs/EBs. (O) Quantification of p-ERK signal in ISC/EBs from midguts as in N. Dashed lines represent a magnified view of cells. Two-way ANOVA followed by Tukey’s multiple comparisons test. * (P<0.1), **** (P<0.0001). Scale bar = 50µm.

### Tracheal derived Pvf1 drives midgut regeneration through activation of dual paracrine and autocrine Pvr/MAPK signaling

We hypothesized that MAPK activating factors other than Bnl/FGF might be regulated by Dh31R signaling in the trachea. Analysis of our TTC Targeted DamID (TaDa) data (Perochon et al., 2021), revealed binding of RNA Pol II to genes encoding for *Drosophila* vascular endothelial growth factor (VEGF) and platelet-derived growth factor (PDGF)-like *Pvf1* and *Pvf3 (Parsons and Foley, 2013)* (Figure S4C). Autocrine Pvf/Pvr signalling drives homeostatic ISC self-renewal through MAPK activation (Bond and Foley, 2012). We hypothesized that tracheal Pvf may activate midgut regeneration by signalling paracrinally to Pvr expressed within ISCs. Interestingly, RNAi knockdown of *Pvf* ligands from TTCs showed that *Pvf1,* but not *Pvf3,* was necessary to induce tracheal remodeling and ISC proliferation during midgut regeneration (Figure 6E-G and S4D). Therefore, the functional role of *Pvf1* produced by TTCs appears distinct from that of tracheal Bnl, which is redundant for TTC remodeling (Perochon et al., 2021). These results suggest that autocrine Pvf1/Pvr signalling within the trachea may impact ISC proliferation indirectly, through the regulation of TTC remodeling. Consistently, tracheal specific knock down of *Pvr* impairs TTC remodeling and MAPK signaling activity and inhibits ISC proliferation in regenerating midguts (Figure 6H-L). Furthermore, knocking down *Pvr* in ISCs/EBs using the *escargot-gal4* driver (*esg^ts^>GFP*) led to almost complete impairment of ISC proliferation and MAPK signalling activity in regenerating midguts (Figure 6M-O).

Altogether, our results uncover a new cellular source and dual signaling mode for *Drosophila* VEGF/PDGF-like ligand, Pvf1, regulating midgut regeneration from the tracheal stem cell niche via ISC and TTC receptors. This distinctive versatile characteristic of tracheal derived Pvf1 makes it a likely candidate for regulation by Dh31/Dh31R signaling during midgut regeneration.

### Dh31R signaling regulates tracheal Pvf1 production in the regenerating midgut

To investigate a potential connection between Dh31/Dh31R signaling and Pvf1, we stained *Dh31R-T2A-Gal4>mCD8-GFP* midguts carrying endogenously tagged Pvf1 (Pvf1-HA) with anti-HA antibody (Figure 7A), which revealed co- localization between the two proteins. Immunostaining to detect either HA-tagged or unmodified Pvf1 revealed strong upregulation of Pvf1 in ECs and TTCs of *Pe* treated midguts (Figure 7B and S5A). Importantly, knocking down *Dh31R* from trachea (Figure 7B) or *Dh31* from neurons (Figure 7D) impaired Pvf1 upregulation in TTCs of the posterior midgut (Figure 7C, E).

**Figure 7:**
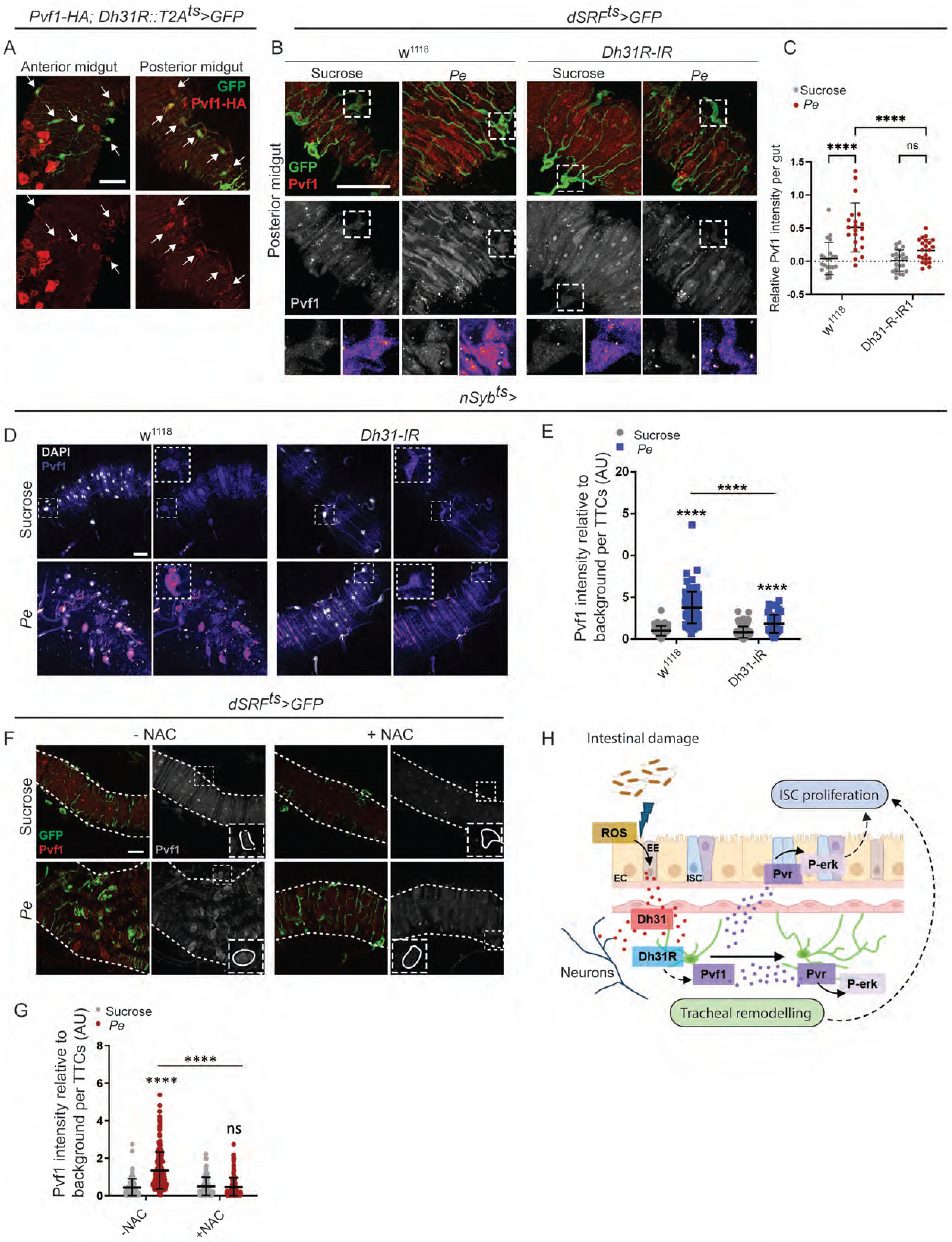
Dh31R signaling regulates tracheal Pvf1 production in the regenerating midgut. (A) Representative images of Pvf1-HA (red) reporter co- localization with GFP driven by *Dh31R-gal4* (green) in the anterior and posterior midgut of animals fed with *Pe*. Arrows indicate Pvf1 (red) co-localization with TTCs (green). (B) Pvf1 immunostaining (red and grey) in posterior midguts upon *Dh31R* KD (*Dh31R-IR1*) in TTCs (green) in animals fed Sucrose or *Pe*. Squares indicate TTCs presented in magnified views in the lowest panels. Magnified cells are colored in white and in fire scale to facilitate visualization. (C) Relative Pvf1 signal intensity in TTCs of midguts as in B. (D) Representative confocal images of Pvf1 (in fire color scale) immunostaining in the posterior midgut upon *Dh31*-KD (*Dh31*-IR) in neurons under the pan-neuronal driver *nSyb-gal4 (nSyb^ts^*) and following treatment with Sucrose or *Pe*. (E) Quantification of Pvf1 signal intensity in TTCs from midguts as in D. (F) Representative confocal images of Pvf1 immunostaining (red and white) in TTCs (green) from posterior midgut of animals fed Sucrose or *Pe* with or without NAC. Squares indicate TTCs presented in magnified views within insets. (G) Quantification of Pvf1 signal intensity levels in TTCs of midguts as in D. (H) Working model. Two-way ANOVA followed by Tukey’s multiple comparisons test. **** (P<0.0001). Dashed lines delineate the edges of the midgut. Scale bar = 50µm.

Tracheal remodeling and Dh31 expression are highly dependent on damage-induced ROS production from the midgut epithelium (Perochon et al., 2021) (Figure 4), we therefore asked whether Pvf1 was also regulated by ROS. Indeed, Pvf1 upregulation was significantly prevented in *Pe* treated animals fed with the antioxidant NAC (Figure 7F, G). Altogether, our results uncover a novel EE/Neuronal/tracheal signaling network controlling global and domain specific ISC proliferation during adult midgut regeneration. Mechanistically, this interorgan signaling system involves regulated production of Dh31 by EE cells in the anterior midgut and constitutive sources of the ligand from EE cells in the posterior midgut and neurons. Dh31 signaling to Dh31R within TTCs induces the production of a previously uncharacterized source of Pvf1 from the trachea. Reciprocally, tracheal Pvf1 induces tracheal remodeling and regenerative ISC proliferation through autocrine and paracrine Pvr/MAPK activation, respectively (Figure 7H).

## Discussion

### Local and global regulation of ISC proliferation by neuroendocrine Dh31 during intestinal regeneration

Our study provides new evidence of complex fine-tuning of intestinal regeneration trough multi-tissue signaling and the importance of damage dependent regulation of EE cell function and specific peptide hormone secretion in this process. Even though EEs have been previously shown to contribute to intestinal homeostasis in flies and mammals (Amcheslavsky et al., 2014, Scopelliti et al., 2014, Drucker et al., 1996, Sasaki et al., 2001), our work represents the first report of EE and neuronal derived peptide hormone and PDGF/VEGF as a vascular stem cell niche regulating intestinal regeneration.

Interestingly, although damage regulated production and secretion of EE Dh31 is specific to the peptide produced in the anterior midgut, the impact of Dh31 on ISC proliferation in the regenerating midgut is global. A similar observation was described in another study, where Dh31 from the anterior midgut is necessary to promote peristaltic muscle contraction throughout the midgut via activation of Dh31-R in the visceral muscle (Benguettat et al., 2018). This could be explained by Dh31 acting not only at short range/paracrinally in the midgut, but also via its release into the extracellular space (Lin et al., 2022) and, therefore, targeting its receptor in regions of the midgut far away from where its production site. Additionally, the 3D conformation of the midgut, which brings the anterior and posterior regions into proximity (Buchon et al., 2013), could facilitate the communication between cells across gut sections.

The area of biggest need for high levels of Dh31 appears to be within the posterior region of the regenerating midgut, where we detected expression and exclusive functionality of an additional source of Dh31, provided by neurons (Figure 2). This may be related to the increased ISC numbers and regenerative activity of the posterior midgut, which is likely to require higher thresholds of proliferative factors and downstream activated signals than other, perhaps more quiescent, sections of the tissue.

Apart from its short-range action within the midgut and midgut associated tissues, Dh31 secretion into the hamolymph has clear systemic consequences (Lin et al., 2022), which we have yet to explore in the context of intestinal damage. EE-derived Dh31 was recently shown to signal to neurons in the brain altering feeding and courtship behaviour (Lin et al., 2022). Dh31-R is highly expressed in neurons in the brain, ventral nerve cord and corpora allata (Deng et al., 2019, Kurogi et al., 2023). Additionally its isoform Dh31-R RC is widely expressed in the glia (Deng et al., 2019). Therefore, damage induced Dh31 from the midgut may regulate systemic traits, which could in turn be potentially instructive to the process of intestinal regeneration and beyond.

We observed a conserved phenomenon of increased EEs proportion during intestinal injury in both flies and mammals. Given the regenerative role of EEs described here and additional protective and immune roles ascribed to this cells (Benguettat et al., 2018, Kamareddine et al., 2018), inducible changes in EE cell numbers might be associated to broad biological functions like worthy of further investigation in flies and mammals.

### Dh31 and its communication with the vascular intestinal stem cell niche

We show here that Dh31 participates in intestinal regeneration via communication with its receptors in the vascular-like tracheal component of the ISC niche. Interestingly, signal relay to associated tissues is a common mechanism used by peptide hormones in both flies as humans (Scopelliti et al., 2014, Amcheslavsky et al., 2014, Estall and Drucker, 2006, Baldassano and Amato, 2014), as perhaps the most efficient strategy to translate local EE cues into global tissue responses. While our results suggest stronger impairment of midgut regeneration when knocking down Dh31 receptor in terminal tracheal cells compared to other niche cells, we cannot rule out a role for Dh31 signaling to receptor in other midgut associated tissues and epithelial cells, as we observed small but significant role in ISC proliferation when affecting Dh31R expression in such cellular compartments.

Dh31 has similarities with the human Calcitonin gene related peptide (CGRP), which is a splicing variant from the Calcitonin gene (Furuya et al., 2000). Similarly, CGRP have very diverse functions in organismal homeostasis, acting locally or systemically in multiple organs. However, CGRP is not found expressed in EE cells but mainly in immune cells and central and peripheral neurons (Russell et al., 2014). Our evidence on the expression and function of neuronal derived Dh31 is a clear parallel to the latter. Indeed, as we observed here, CGRP communicates with the cardiovascular system and is known to be a potent vasodilator by activating cAMP pathway through its receptor in endothelial cells and the smooth muscle that surrounds them (Russell et al., 2014). Additionally, CGRP has also been reported to have anti and pro-inflammatory roles and facilitates wound healing possibly by local upregulation of growth factors such as VEGF, FGF, TGF-β (Toda et al., 2008, Mishima et al., 2011). In the intestine, new studies are beginning to show the importance of CGRP in inflammatory contexts (Xu et al., 2019, Manion et al., 2023). However, its CGRP functions in tissue regeneration are largely unexplored and our study provides the first evidence of such a role in the fly midgut, prompting similar investigations in mammals.

## Materials and Methods

### Fly stocks and husbandry

A complete list of fly strains used in this paper is included in the Key Resources Table. All flies were kept in temperature-controlled incubators with a 12-12-hour light-dark cycle on cornmeal-based rearing medium. Fly stocks were kept at 18°C. Crosses for experiments containing the temperature sensitive *tubulin*-Gal80^ts^ were set up at 18°C. Their progeny was kept for 3 days in fresh standard medium in the same temperature as parental crosses and then transferred to 29°C for 5-8 days, unless otherwise stated. Flies at 29°C were transferred to fresh medium every two days. Other crosses were set up and kept at 25°C. Only mated female flies were used for experiments.

### Intestinal damage assays

10 days old mated female flies were fed a 5% sucrose solution (Mock) or sucrose containing *Pseudomonas entomophila* (*Pe*) at an OD_600_ = 50 for 16h, 50µg/ml bleomycin (Sigma-Aldrich, cat no. B2434) for 24h or 3% DSS (Sigma-Aldrich, cat no. 42867) for 48h applied on a glass microfiber filter (Whatman). Fresh solution containing DSS was applied daily. Oxidative stress was induced by feeding 1% hydrogen peroxide (H_2_O_2_) diluted in 5% sucrose for 16h.

### NAC treatment

Prior to intestinal damage generation, flies were fed a 5% sucrose solution containing 20mM NAC (Sigma-Aldrich, cat no. A7250) for 24h. Then flies were fed either 5% sucrose + NAC or 5% sucrose + NAC + *Pe* (OD_600_ = 50) for an additional period of 16h.

### Capsaicin treatment

Flies were fed a 5% sucrose solution (Mock) or sucrose containing capsaicin (50µM) for 16h applied on a glass microfiber filter.

### Immunofluorescence

Guts from adult female flies were dissected in PBS and immediately fixed in 4% formaldehyde (FA, EM grade, Polysciences) diluted in PBS for 1h at room temperature (RT) and then washed three times in PBS-T (PBS + 0.2% Triton X100). Samples were incubated with the primary antibody diluted in PBT (PBS-T + 2 % BSA) overnight at 4°C. Guts were washed three times in PBS-T and then incubated with secondary antibody in PBT for 2h at RT. After incubation, guts were washed three times for 20min in PBS-T and mounted on glass slides with 13mm x 0.12mm spacers (sigma) in Vectashield mounting medium with DAPI (Vector Laboratories). Guts stained with anti-DH31, were blocked in 7% goat serum (in PBS-T) for 1h before primary antibody incubation. For anti-PVF1 and anti-pMAPK staining, fixation was performed in 4% formaldehyde for 1h, followed by 5min methanol fixation added to the sample gradually and then 10min in 100% methanol. Samples were blocked in 7% goat serum for 1h before primary antibody incubation.

For brains and ventral nerve cord (VNC) dissection, whole flies or fly heads were first washed in 70% ethanol for 30sec and fixed as described above. Heads were transferred to PBS-T and dissected. After, they were washed two times in PBS-T and blocked in 7% goat serum for at least 1h. Samples were incubated with primary antibodies in PBT for 48 hours at 4°C followed by the steps mentioned above.

A list of all antibodies and dilutions used in this study is included in the Key Resources Table.

### Tissue imaging

Confocal microscope images were taken on a Zeiss LSM 710, Zeiss LSM 780 or Zeiss LSM 880 using identical acquisition conditions for all samples from a given experiment. Images were processed on ImageJ and Photoshop.

Alternatively, Images were taken with the automated confocal microscope Opera Phenix (Perkin Elmer) using a 5X objective for pre-scan and 20X water objective for re-scan. Images were processed and analysed with Harmony (Perkin Elmer) or ImageJ.

### Quantifications from immunofluorescence images

Confocal images from the anterior and/or posterior midgut were used to quantify signal intensity using ImageJ. PVF1 signal was obtained from maximal projections of a limited number of Z-stacks for each terminal trachea cell. For Dh31 quantifications, whole gut images from the Opera Phenix were exported as Tiff files and stitched. Regions of interest were cropped and quantified with ImageJ using a MACRO (Appendix 1).

Enteroendocrine cell proportion was obtained from whole midgut images acquired with Opera Phenix. Total number of Prospero+ objects were divided by the total number of DAPI+ objects using Harmony (Perkin Elmer).

Total number of pH3+ cells were scored manually on visual inspection using an epifluorescent microscope Olympus BX51 in the anterior and/or posterior midgut.

### Terminal tracheal cells coverage

Whole posterior midgut images were dragged to ImageJ and only the green channel exhibiting TTCs labelled by GFP was analysed. Maximal projection of all stacks had their threshold adjusted without creating a noisy background. Images were then skeletonized and the exact area to be quantified was selected. Measurement of signal coverage in a specific area was analysed.

### Analysis of calcium levels in Dh31+ cells

Crosses and rearing of *UAS-GCaMP7s* and *Dh31-Gal4* were done at 25°C. 5-8 days old mated female flies expressing GCaMP7s specifically in Dh31+ cells were treated with 5% sucrose solution (Mock) or sucrose containing *Pe* at an OD_600_ = 50 for 16h on a glass microfiber filter. Whole guts were carefully dissected in Schneider’s Drosophila-Medium (Gibco) and directly mounted on slides containing the same medium. Guts were immediately observed with a Zeiss Observer 7. The number of green fluorescent cells was scored manually by visual inspection in the anterior and posterior midgut.

### RNA extraction and RT-qPCR

15 midguts, 30 anterior/posterior midguts or 30 heads were transferred immediately to 800 µL TRIzol (Ambion by Life technologies), smashed with a pestle and stored at -80°C until RNA extraction. RNA was extracted by precipitation. cDNA was obtained using the high-capacity cDNA reverse transcription kit (Applied Biosystems) following manufacturers protocol. Quantitative PCR was done at least in biological triplicates for each genotype/condition and run in technical triplicates on an Applied Biosystems 7500 Fast Real-Time PCR machine using PerfeCta SYBR Green FastMix (Quanta). Data was analysed using QuantStudio design and analysis desktop software v1.4.3. Expression of target genes was normalized to the internal control Rpl32 using standard curves. A list of primer sequences can be found in the Key Resources Table.

### Mouse intestinal regeneration and immunohistochemistry

Mice (*Mus musculus*) were crossed to a C57BL/6 background and were subjected to 10Gy- 72h before the experiment. For colitis induction, mice were fed a 2% solution of DSS in water and weighed daily. They were fed 2% DSS for 5 days and allowed to rest for 2 days. Any mouse that lost >20% of its body weight or exhibiting clinical signs such as hunching or excessive diarrhoea was humanely culled. Mice were then humanely killed by a rising concentration of CO2. The small intestine and colon were isolated and flushed with tap water. 10 1-cm portions of intestine or colon were bound together with surgical tape and fixed in 10% neutral buffered formalin between 20 to 48h and then transferred to 70% ethanol and paraffinized. Paraffinized tissue were cut in 4µm sections and placed onto slides and incubated at 60°C overnight. Before staining, the sections were dewaxed for 5min in xylene, followed by rehydration through decreasing concentrations of alcohol and a final wash with H_2_O for 5min. The formalin-fixed paraffin-embedded sections underwent heat-induced epitope retrieval in a Dako pretreatment module. The sections were heated in Target retrieval solution high pH (Dako, cat. no. K8004) for 20min at 97 °C before cooling to 65 °C. The slides were removed and washed in Tris-buffered saline with Tween (TbT; Dako, cat. no. K8000) and loaded onto a Dako autostainer link 48 platform where they were stained with anti- Chromogranin A antibody (1:700; Proteintech, cat. no. 23342-1-AP) following standard immunohistochemistry procedures and stained with Haematoxylin for 5min. All experiments were performed in accordance with UK Home Office regulations which undergoes local ethical review at Glasgow University.

## Key Resources Table

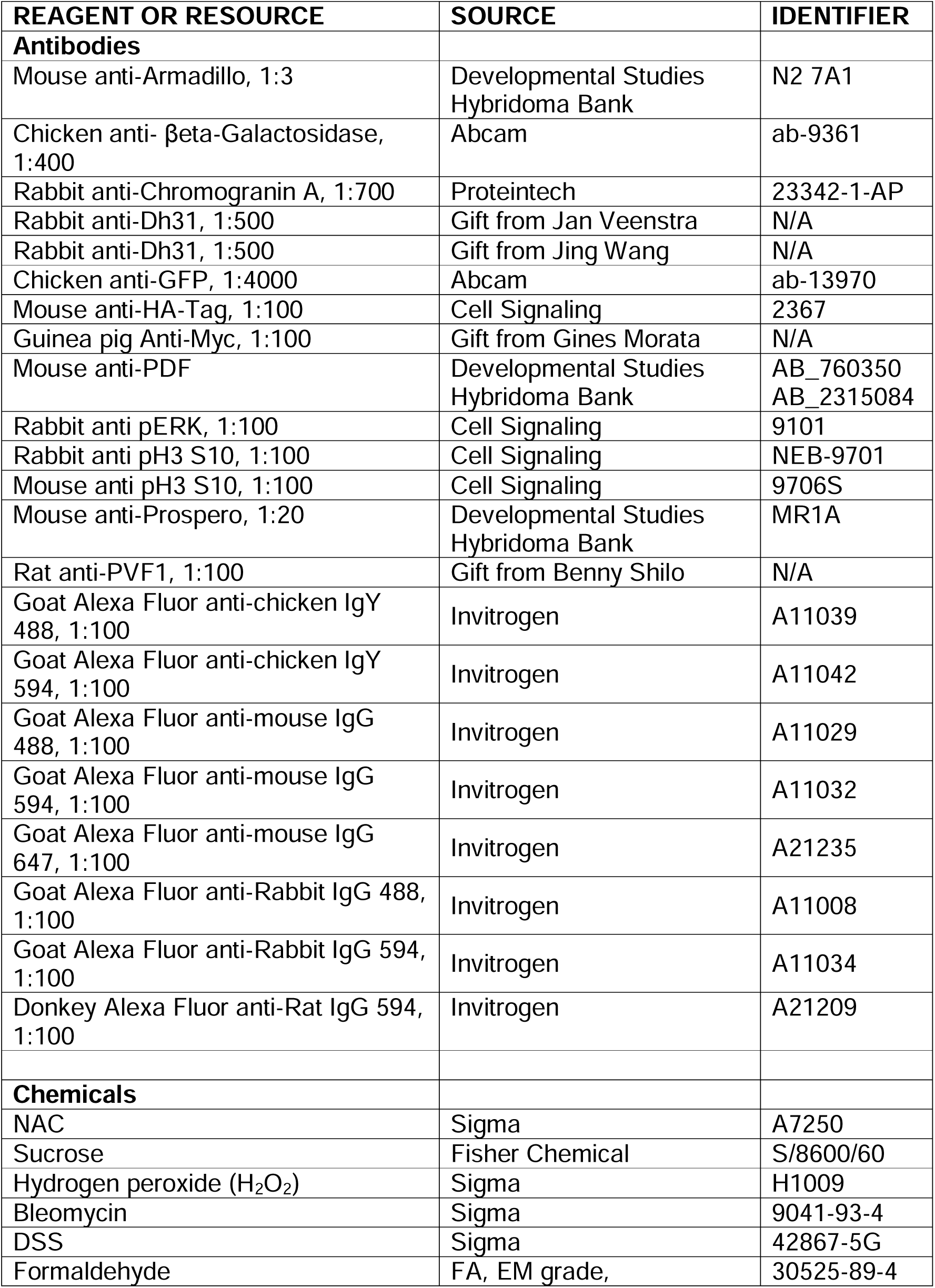
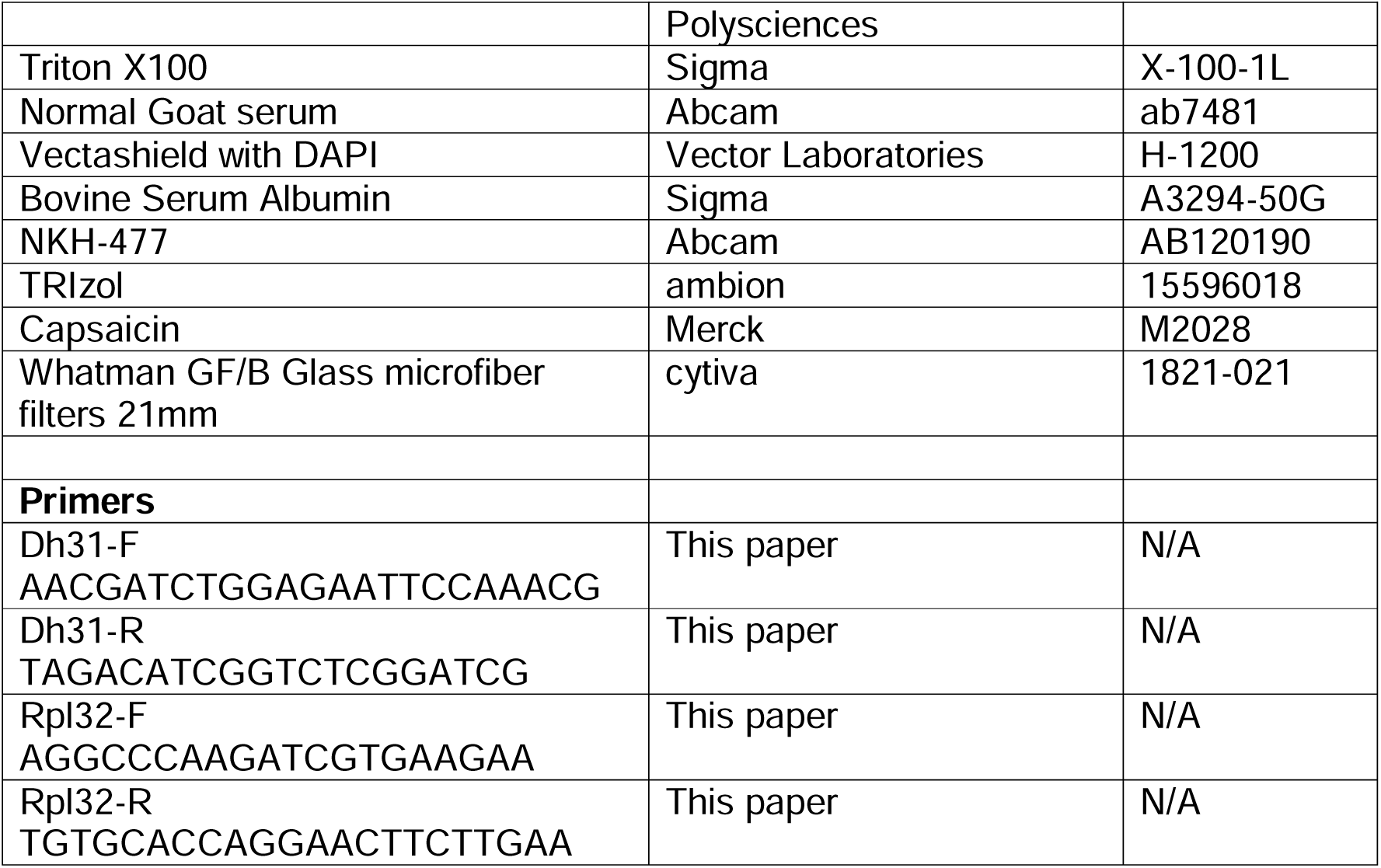

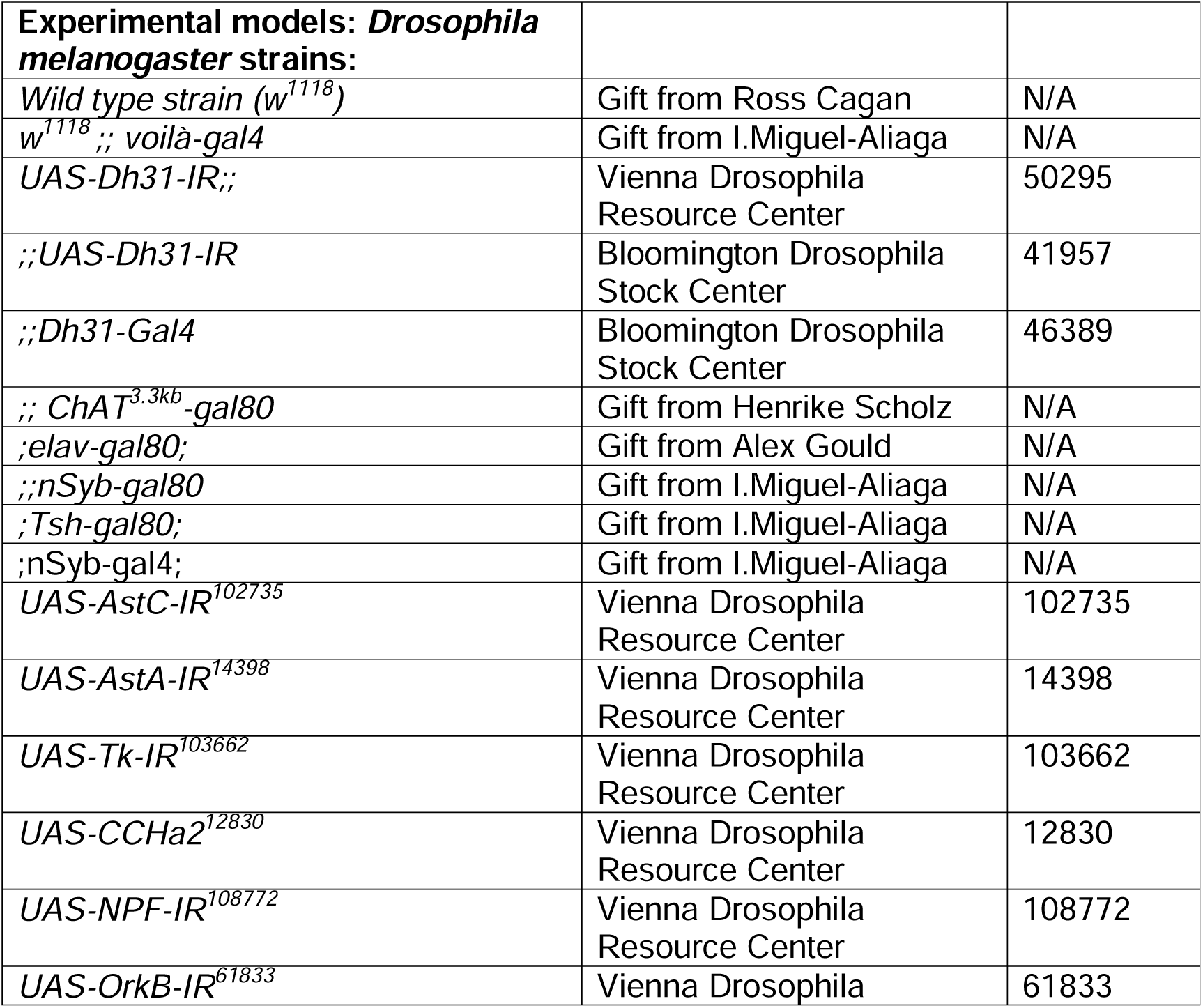
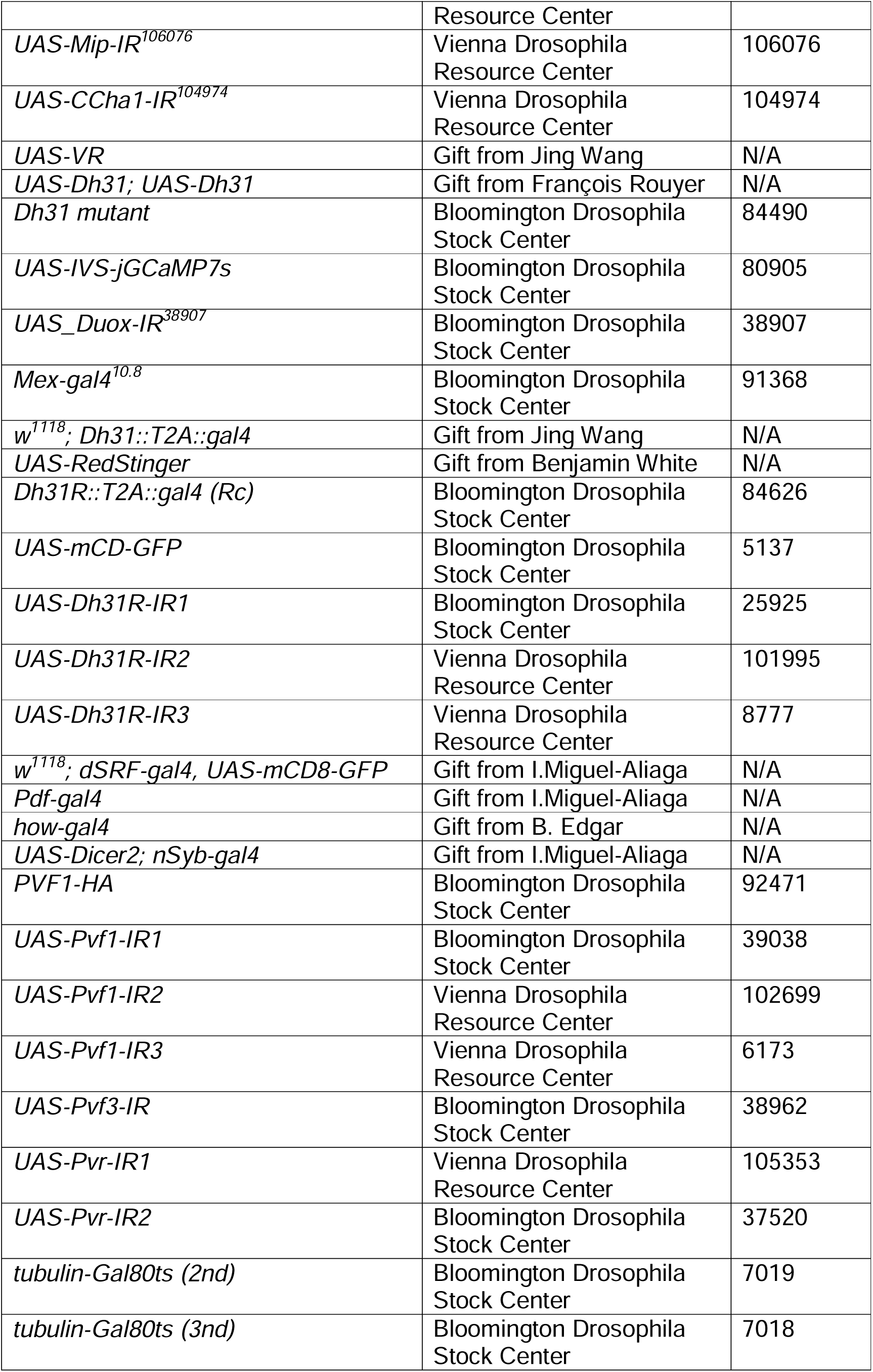
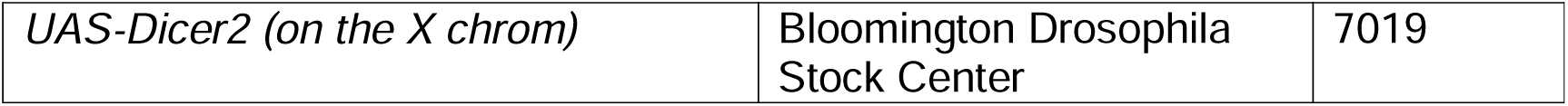

## Detailed genotypes

Figure 1:

1. *_w_1118*
2. *w[*]; TI{RFP[3xP3.cUa]=TI}Dh31[attP] (Dh31 knock out)*
3. *UAS-Dicer2; tub-gal80^ts^; voilà-gal4*
4. *UAS-Dicer2/UAS-Dh31-IR^50295^; tub-gal80^ts^; voilà-gal4*
5. *UAS-Dicer2; tub-gal80^ts^; UAS-Dh31-IR^41957^/ voilà-gal4*
6. *w^1118^; tub-gal80^ts^/ Dh31-gal4*
7. *UAS-Dh31-IR^50295^; tub-gal80^ts^/ Dh31-gal4*
8. *w^1118^; tub-gal80^ts^/ UAS-Dh31-IR^41957^/ Dh31-gal4*
9. *UAS-Dicer2; tub-gal80^ts^; voilà-gal4/ ChAT^3.3kb^-gal80*
10. *UAS-Dicer2; tub-gal80^ts^; UAS-Dh31-IR^41957^/voilà-gal4/ ChAT^3.3kb^-gal80*
11. *w^1118^; ;UAS-Dh31-IR^41957^*

Figure 2

1. UAS-Dicer2; nSyb-gal4/ UAS-GFP; tub-gal80^ts^
2. *UAS-Dicer2; nSyb-gal4;/UAS-GFP; UAS-Dh31-IR^41957^/ tub-gal80^ts^*
3. *w^1118^; ;UAS-Dh31-IR^41957^*

Figure 3

1. w^1118^; tub-gal80^ts^; Dh31-gal4
2. *_w_1118*
3. *w^1118^; ;Dh31-gal4*
4. *w^1118^; UAS-IVS-jGCaMP7s; Dh31-gal4*

Figure 4

1. _w_1118
2. *w^1118^; Mex-gal4^10.8^; tub-gal80^ts^*
3. *w^1118^; UAS-Duox-IR^38907^/ Mex-gal4^10.8^; tub-gal80^ts^*
4. *w^1118^; tub-gal80^ts^; Dh31-gal4*

Figure 5

1. w^1118^; Dh31R::T2A::gal4 (Rc)/ UAS-mCD8-GFP;
2. *w^1118^; dSRF-gal4, UAS-mCD8-GFP; tub-gal80^ts^*
3. *w^1118^; dSRF-gal4, UAS-mCD8-GFP; tub-gal80^ts^/ UAS-Dh31R-IR^25925^*
4. *w^1118^; dSRF-gal4, UAS-mCD8-GFP / UAS-Dh31R-IR^101995^; tub-gal80^ts^*
5. *w^1118^; dSRF-gal4, UAS-mCD8-GFP / UAS-Dh31R-IR^8777^; tub-gal80^ts^*

Figure 6

1. *w^1118^; dSRF-gal4, UAS-mCD8-GFP; tub-gal80^ts^*
2. *w^1118^; dSRF-gal4, UAS-mCD8-GFP; tub-gal80^ts^/ UAS-Dh31R-IR^25925^*
3. *w^1118^; dSRF-gal4, UAS-mCD8-GFP / UAS-Dh31R-IR^101995^; tub-gal80^ts^*
4. *w^1118^; dSRF-gal4, UAS-mCD8-GFP/ UAS-Pvf1^39038^-IR; tub-gal80^ts^*
5. *w^1118^; dSRF-gal4, UAS-mCD8-GFP/UAS-Pvf1-IR^102699;^ tub-gal80^ts^*
6. *w^1118^; dSRF-gal4, UAS-mCD8-GFP/ UAS-Pvr-IR^105353^; tub-gal80^ts^*
7. *w^1118^; dSRF-gal4, UAS-mCD8-GFP/UAS-Pvr-IR^37520^; tub-gal80^ts^*
8. *w^1118^; esg-gal4, UAS-mCD8-GFP; tub-gal80^ts^*
9. *w^1118^; UAS-Pvr-IR^105353^;*
10. *w^1118^; esg-gal4, UAS-mCD8-GFP/UAS-Pvr-IR^105353^; tub-gal80^ts^*

Figure 7

1. *Pvf1-HA; Dh31R::T2A-gal4/UAS-mCD8-GFP; tub-gal80^ts^*
2. *w^1118^; dSRF-gal4, UAS-mCD8-GFP; tub-gal80^ts^*
3. *w^1118^; dSRF-gal4, UAS-mCD8-GFP; tub-gal80^ts^/ UAS-Dh31R-IR^25925^*
4. *UAS-Dicer2; nSyb-gal4/ UAS-GFP; tub-gal80^ts^*
5. *UAS-Dicer2; nSyb-gal4;/UAS-GFP; UAS-Dh31-IR^41957^/ tub-gal80^ts^*

Supplementary Figure 1

1. *UAS-Dicer2; tub-gal80^ts^; voilà-gal4*
2. *UAS-Dicer2; UAS-AstC-IR^102735^/ tub-gal80^ts^; voilà-gal4*
3. *UAS-Dicer2; UAS-AstA-IR^14398^/ tub-gal80^ts^; voilà-gal4*
4. *UAS-Dicer2; UAS-Tk-IR^103662^/ tub-gal80^ts^; voilà-gal4*
5. *UAS-Dicer2/UAS-Dh31-IR^50295^; tub-gal80^ts^; / voilà-gal4*
6. *UAS-Dicer2; tub-gal80^ts^; UAS-CCHa2^12830^/ voilà-gal4*
7. *UAS-Dicer2; UAS-NPF-IR^108772^/ tub-gal80^ts^; voilà-gal4*
8. *UAS-Dicer2; UAS-OrkB-IR^61833^/ tub-gal80^ts^; voilà-gal4*
9. *UAS-Dicer2; UAS-Mip-IR^106076^/ tub-gal80^ts^; voilà-gal4*
10. *UAS-Dicer2; tub-gal80^ts^; UAS-CCha1-IR*^104974^*/ voilà-gal4*
11. *UAS-Dicer2; tub-gal80^ts^; voilà-gal4/ ChAT^3.3kb^-gal80*
12. *UAS-Dicer2; tub-gal80^ts^; UAS-Dh31-IR^41957^/voilà-gal4/ ChAT^3.3kb^-gal80*
13. *UAS-Dicer2; tub-gal80^ts^; voilà-gal4/ ChAT^3.3kb^-gal80*
14. *UAS-Dicer2; tub-gal80^ts^/ elav-gal80; voilà-gal4/*
15. *UAS-Dicer2; tub-gal80^ts^; voilà-gal4/ nSyb-gal80*
16. *UAS-Dicer2; tub-gal80^ts^/ tsh-gal80; voilà-gal4/*
17. *w^1118^; tub-gal80^ts^;Dh31-gal4*
18. *w^1118^; UAS-VR/ tub-gal80^ts^/ Dh31-gal4*
19. *w^1118^; UAS-VR/ tub-gal80^ts^/ UAS-Dh31/ Dh31-gal4*

Supplementary Figure 2

1. *w^1118^; UAS-mCD8-GFP; Dh31-gal4*
2. *w^1118^; Dh31-T2A-gal4/ UAS-IVS-jGCaMP7s; UAS-redStinger*

Supplementary Figure 3

1. *w^1118^; Dh31R::T2A::gal4 (Rc)/ UAS-mCD8-GFP;*
2. *w^1118^; Pdf-gal4; tub-gal80^ts^*
3. *w^1118^; Pdf-gal4; tub-gal80^ts^/ UAS-Dh31R-IR^25925^*
4. *UAS-Dicer2; tub-gal80ts; how-gal4*
5. *UAS-Dicer2; tub-gal80ts; how-gal4/ UAS-Dh31R-IR^25925^*
6. *w^1118^; Mex-gal4^10.8^; tub-gal80^ts^*
7. *w^1118^; Mex-gal4^10.8^; tub-gal80^ts^/ UAS-Dh31R-IR^25925^*
8. *w^1118^; dSRF-gal4, UAS-mCD8-GFP; tub-gal80^ts^*
9. *w^1118^; ; UAS-Dh31R-IR^25925^*
10. *w^1118^; dSRF-gal4, UAS-mCD8-GFP; tub-gal80^ts^/ UAS-Dh31R-IR^25925^*
11. *w^1118^; UAS-Dh31R-IR^101995^;*
12. *w^1118^; dSRF-gal4, UAS-mCD8-GFP / UAS-Dh31R-IR^101995^; tub-gal80^ts^*

Supplementary Figure 4

1. *w^1118^; dSRF-gal4, UAS-mCD-GFP; tub-gal80^ts^/bnl-LacZ*
2. *w^1118^; dSRF-gal4, UAS-mCD-GFP/UAS-Dh31R-IR*^101995^*; tub-gal80^ts^/bnl-LacZ*
3. *w^1118^; dSRF-gal4, UAS-mCD-GFP; tub-gal80^ts^*
4. *w^1118^; dSRF-gal4, UAS-mCD-GFP/ UAS-Pvf3^38962^-IR; tub-gal80^ts^*
5. *w^1118^; dSRF-gal4, UAS-mCD-GFP/UAS-Pvr-IR^105353^; tub-gal80^ts^*

Supplementary Figure 5

1. *Pvf1-HA; dSRF-gal4, UAS-mCD-GFP; tub-gal80^ts^*

**Supplementary Figure 1:** (A, B) Quantification of PH3+ cells in the posterior midgut of animals treated with Sucrose or *Pe* following different peptide hormones knockdown in EEs. (C) Representative confocal images of GFP expression (green) in the Brain, VNC and midgut driven by *voilá-gal4* in combination with different neuronal *gal80* lines. EE cells in midguts are stained with anti-Prospero (Pros; red). (D) Dh31 immunostaining (green) in the ventral nerve cord (VNC) and anterior midgut upon *Dh31* KD (*Dh31-IR2*) using the gut specific line (*voilá-gal4* with *ChAT- gal80)*. Nc82 (neuronal marker, magenta). (E) Representative images of Dh31 immunostaining (red and grey) upon overexpression of VR in Dh31+ cells and treatment with its ligand Capsaicin. (F) Quantification of PH3+ cells in control midguts, midguts overexpressing *VR* only or *VR* and *Dh31* in Dh31+ cells following exposure to Sucrose or Capsaicin. (A, B, E) Two-way ANOVA followed by Tukey’s multiple comparisons test. * (P<0,04), ** (P<0,008), *** (P<0.001), **** (P<0.0001). Scale bar = 50µm.

**Supplementary Figure 2:** (A) Dh31 expression pattern (grey) in the midgut labeled with anti-Dh31 (top) or GFP driven by *Dh31-gal4* (bottom). (B) Co- localization of GCamp7 (green) and RedStinger (red) driven by *Dh31* knock in *gal4* in animals fed Sucrose or *Pe* for 16h. Dashed delineate the edges of the midgut. Scale bar = 50µm.

**Supplementary Figure 3:** (A) Expression pattern of *Dh31R* knock-in *gal4* visualized by GFP (green) in enteric neurons, visceral muscle and enterocytes. (B) Co-localization of GFP (green) in enteric neurons with anti-Pdf (grey). (C) Quantification of PH3+ cells in the posterior midgut of animals treated with Sucrose or *Pe* following *Dh31R* knockdown (*Dh31R-*IR1) in Pdf+ enteric neurons (C), visceral muscle (D) and enterocytes (E). (F) Quantification of PH3+ cells in the posterior midgut of animals treated with Sucrose or *Pe* following *Dh31R* knockdown (*Dh31R-* IR1, *Dh31R-*IR1) in TTCs and including UAS-*Dh31R-IR* crossed with *w^1118^* as an additional control. Two-way ANOVA followed by Tukey’s multiple comparisons test. * (P<0,03), ** (P<0,009), **** (P<0.0001). Scale bar = 100µm.

**Supplementary Figure 4:** (A) Representative confocal images of *bnl-LacZ* reporter immunostaining (red and grey) in the posterior midgut upon *Dh31R* KD (*Dh31R-*IR1) in TTCs (green) after treatment with Sucrose or *Pe.* Arrows indicate TTCs. (B) Quantification of *bnl-LacZ* signal intensity values in TTCs from midguts as in A. Grey dots represent single cell values. Black or red dots represent average per midgut. (C) Profile of of RNA Polymerase II binding on *Pvf1* and *Pvf3* gene loci obtained from a published TaDa dataset from TTCs. (D) Quantification of PH3+ cells and tracheal coverage in the posterior midgut upon *Pvf3* KD in TTCs after treatment with Sucrose or *Pe*. (E) Representative confocal images of tracheal coverage (green) in the posterior midgut upon *Pvr* KD (*Pvr*-IR1) in animals treated with Sucrose or *Pe*. Two-way ANOVA followed by Tukey’s multiple comparisons test. ** (P<0.003), **** (P<0.0001). Scale bar = 50µm.

**Supplementary Figure 5:** (A) Pvf1-HA reporter levels in anterior and posterior midguts treated with Sucrose or *Pe.* TTCs (green), Pvf1 (red/white). Squares indicate TTCs shown in magnified views in the lowest panels. (B) Quantification of Pvf1-HA in TTCs of midguts as in A. Scale bar = 25µm.

## Acknowledgements

We would like to thank Irene Miguel-Aliaga, Henrike Scholz, Jan Veentra, Benny Shilo and Jing W. Wang for generously sharing flies and reagents. We thank the Bloomington Drosophila Stock Centre; the Vienna Drosophila Resource Center and the *Drosophila* Studies Hybridoma Bank for fly stocks and antibodies. We would like to thank the Core Services and Advanced Technologies at the CRUK Scotland Institute, which is core funded by Cancer Research UK (A31287). With particular thanks to Leo Carlin, Claire Mitchell, Lynn McGarry and Peter Thomason from the Beatson Advanced Imaging Resource, and Colin Nixon and the Histology Laboratory for assistance with mouse sample preparation and staining. We thank all members of the Cordero laboratory for general advice on the project.

## References

Amcheslavsky, A., Jiang, J. & Ip, Y. T. 2009. Tissue damage-induced intestinal stem cell division in Drosophila. Cell Stem Cell, 4, 49–61.

Amcheslavsky, A., Song, W., Li, Q., Nie, Y., Bragatto, I., Ferrandon, D., Perrimon, N. & Ip, Y. T. 2014. Enteroendocrine cells support intestinal stem-cell-mediated homeostasis in Drosophila. Cell Rep, 9, 32–39.

Baldassano, S. & Amato, A. 2014. GLP-2: What do we know? What are we going to discover? Regulatory Peptides, 194–195, 6-10.

Benguettat, O., Jneid, R., Soltys, J., Loudhaief, R., Brun-Barale, A., Osman, D. & Gallet, A. 2018. The DH31/CGRP enteroendocrine peptide triggers intestinal contractions favoring the elimination of opportunistic bacteria. PLOS Pathogens, 14, e1007279.

Beumer, J., Puschhof, J., Bauzá-Martinez, J., Martínez-Silgado, A., Elmentaite, R., James, K. R., Ross, A., Hendriks, D., Artegiani, B., Busslinger, G. A., Ponsioen, B., Andersson-Rolf, A., Saftien, A., Boot, C., Kretzschmar, K., Geurts, M. H., BAR-Ephraim, Y. E., Pleguezuelos-Manzano, C., Post, Y., Begthel, H., Van Der Linden, F., Lopez-Iglesias, C., Van De Wetering, W. J., Van Der Linden, R., Peters, P. J., Heck, A. J. R., Goedhart, J., Snippert, H., Zilbauer, M., Teichmann, S. A., Wu, W. & Clevers, H. 2020. High-Resolution mRNA and Secretome Atlas of Human Enteroendocrine Cells. Cell, 181, 1291–1306.e19.

Bond, D. & Foley, E. 2012. Autocrine platelet-derived growth factor-vascular endothelial growth factor receptor-related (Pvr) pathway activity controls intestinal stem cell proliferation in the adult Drosophila midgut. J Biol Chem, 287, 27359–70.

Bosi, G., Shinn, A. P., Giari, L., Simoni, E., Pironi, F. & Dezfuli, B. S. 2005. Changes in the neuromodulators of the diffuse endocrine system of the alimentary canal of farmed rainbow trout, Oncorhynchus mykiss (Walbaum), naturally infected with Eubothrium crassum (Cestoda). J Fish Dis, 28, 703–11.

Buchon, N., Osman, D., David, F. P., Fang, H. Y., Boquete, J. P., Deplancke, B. & Lemaitre, B. 2013. Morphological and molecular characterization of adult midgut compartmentalization in Drosophila. Cell Rep, 3, 1725–38.

Chen, J., Xu, N., Wang, C., Huang, P., Huang, H., Jin, Z., Yu, Z., Cai, T., Jiao, R. & Xi, R. 2018. Transient Scute activation via a self-stimulatory loop directs enteroendocrine cell pair specification from self-renewing intestinal stem cells. Nat Cell Biol, 20, 152–161.

Cognigni, P., Bailey, A. P. & Miguel-Aliaga, I. 2011. Enteric Neurons and Systemic Signals Couple Nutritional and Reproductive Status with Intestinal Homeostasis. Cell Metabolism, 13, 92–104.

Dahly, E. M., Gillingham, M. B., Guo, Z., Murali, S. G., Nelson, D. W., Holst, J. J. & Ney, D. M. 2003. Role of luminal nutrients and endogenous GLP-2 in intestinal adaptation to mid-small bowel resection. Am J Physiol Gastrointest Liver Physiol, 284, G670–82.

Deng, B., Li, Q., Liu, X., Cao, Y., Li, B., Qian, Y., Xu, R., Mao, R., Zhou, E., Zhang, W., Huang, J. & Rao, Y. 2019. Chemoconnectomics: Mapping Chemical Transmission in Drosophila. Neuron, 101, 876–893.e4.

Drucker, D. J., Erlich, P., Asa, S. L. & Brubaker, P. L. 1996. Induction of intestinal epithelial proliferation by glucagon-like peptide 2. Proc Natl Acad Sci U S A, 93, 7911–6.

Dutta, D., Dobson, ADAM J., Houtz, PHILIP L., Gläßer, C., Revah, J., Korzelius, J., Patel, P. H., Edgar, BRUCE A. & Buchon, N. 2015. Regional Cell-Specific Transcriptome Mapping Reveals Regulatory Complexity in the Adult Drosophila Midgut. Cell Reports, 12, 346–358.

Estall, J. L. & Drucker, D. J. 2006. Glucagon-like Peptide-2. Annu Rev Nutr, 26, 391–411.

Forbes, A. B., Warren, M., Upjohn, M., Jackson, B., Jones, J., Charlier, J. & Fox, M. T. 2009. Associations between blood gastrin, ghrelin, leptin, pepsinogen and Ostertagia ostertagi antibody concentrations and voluntary feed intake in calves exposed to a trickle infection with O. ostertagi. Vet Parasitol, 162, 295–305.

Furuya, K., Milchak, R. J., Schegg, K. M., Zhang, J., Tobe, S. S., Coast, G. M. & Schooley, D. A. 2000. Cockroach diuretic hormones: Characterization of a calcitonin-like peptide in insects. Proceedings of the National Academy of Sciences, 97, 6469–6474.

Gehart, H., VAN Es, J. H., Hamer, K., Beumer, J., Kretzschmar, K., Dekkers, J. F., Rios, A. & Clevers, H. 2019. Identification of Enteroendocrine Regulators by Real-Time Single-Cell Differentiation Mapping. Cell, 176, 1158–1173.e16.

Gribble, F. M. & Reimann, F. 2016. Enteroendocrine Cells: Chemosensors in the Intestinal Epithelium. Annual Review of Physiology, 78, 277–299.

Guo, X., Lv, J. & Xi, R. 2022. The specification and function of enteroendocrine cells in Drosophila and mammals: a comparative review. Febs j, 289, 4773–4796.

Guo, X., Yin, C., Yang, F., Zhang, Y., Huang, H., Wang, J., Deng, B., Cai, T., Rao, Y. & Xi, R. 2019. The Cellular Diversity and Transcription Factor Code of Drosophila Enteroendocrine Cells. Cell Reports, 29, 4172–4185.e5.

Hageman, J. H., Heinz, M. C., Kretzschmar, K., Van Der Vaart, J., Clevers, H. & Snippert, H. J. G. 2020. Intestinal Regeneration: Regulation by the Microenvironment. Developmental Cell, 54, 435–446.

Harnack, C., Berger, H., Antanaviciute, A., Vidal, R., Sauer, S., Simmons, A., Meyer, T. F. & Sigal, M. 2019. R-spondin 3 promotes stem cell recovery and epithelial regeneration in the colon. Nat Commun, 10, 4368.

Harrison, E., Lal, S. & Mclaughlin, J. T. 2013. Enteroendocrine cells in gastrointestinal pathophysiology. Curr Opin Pharmacol, 13, 941–5.

He, L., Si, G., Huang, J., Samuel, A. D. T. & Perrimon, N. 2018. Mechanical regulation of stem-cell differentiation by the stretch-activated Piezo channel. Nature, 555, 103–106.

Hung, R. J., Hu, Y., Kirchner, R., Liu, Y., Xu, C., Comjean, A., Tattikota, S. G., Li, F., Song, W., HO Sui, S. & Perrimon, N. 2020. A cell atlas of the adult Drosophila midgut. Proc Natl Acad Sci U S A, 117, 1514–1523.

Jiang, H., Grenley, M. O., Bravo, M.-J., Blumhagen, R. Z. & Edgar, B. A. 2011. EGFR/Ras/MAPK Signaling Mediates Adult Midgut Epithelial Homeostasis and Regeneration in <EM>Drosophila</EM>. Cell Stem Cell, 8, 84–95.

Jiang, H., Patel, P. H., Kohlmaier, A., Grenley, M. O., Mcewen, D. G. & Edgar, B. A. 2009a. Cytokine/Jak/Stat Signaling Mediates Regeneration and Homeostasis in the <em>Drosophila</em> Midgut. Cell, 137, 1343–1355.

Jiang, H., Patel, P. H., Kohlmaier, A., Grenley, M. O., Mcewen, D. G. & Edgar, B. A. 2009b.Cytokine/Jak/Stat signaling mediates regeneration and homeostasis in the Drosophila midgut. Cell, 137, 1343–55.

Kamareddine, L., Robins, W. P., Berkey, C. D., Mekalanos, J. J. & Watnick, P. I. 2018. The Drosophila Immune Deficiency Pathway Modulates Enteroendocrine Function and Host Metabolism. Cell Metab, 28, 449–462.e5.

Kim, J.-E., Fei, L., Yin, W.-C., Coquenlorge, S.,Rao-Bhatia, A., Zhang, X., Shi, S. S. W., Lee, J. H., Hahn, N. A., Rizvi, W., Kim, K.-H., Sung, H.-K., Hui, C.-C., Guo, G. & Kim, T.-H. 2020. Single cell and genetic analyses reveal conserved populations and signaling mechanisms of gastrointestinal stromal niches. Nature Communications, 11, 334.

Kitamoto, T. 2002. Conditional disruption of synaptic transmission induces male-male courtship behavior in Drosophila. Proc Natl Acad Sci U S A, 99, 13232–7.

Kurogi, Y., Imura, E., Mizuno, Y., Hoshino, R., Nouzova, M., Matsuyama, S., Mizoguchi, A., Kondo, S., Tanimoto, H., Noriega, F. G. & Niwa, R. 2023. Female reproductive dormancy in Drosophila is regulated by DH31-producing neurons projecting into the corpus allatum. Development, 150.

Lebrun, L. J., Lenaerts, K., Kiers, D., PAIS DE Barros, J. P., LE Guern, N., Plesnik, J., Thomas, C., Bourgeois, T., Dejong, C. H. C., Kox, M., Hundscheid, I. H. R., Khan, N. A., Mandard, S., Deckert, V., Pickkers, P., Drucker, D. J., Lagrost, L. & Grober, J. 2017. Enteroendocrine L Cells Sense LPS after Gut Barrier Injury to Enhance GLP-1 Secretion. Cell Rep, 21, 1160–1168.

Li, H., Janssens, J., DE Waegeneer, M., Kolluru, S. S., Davie, K., Gardeux, V., Saelens, W., David, F. P. A., Brbić, M., Spanier, K., Leskovec, J., Mclaughlin, C. N., Xie, Q., Jones, R. C., Brueckner, K., Shim, J., Tattikota, S. G., Schnorrer, F., Rust, K., Nystul, T. G., CARVALHO-Santos, Z., Ribeiro, C., Pal, S., Mahadevaraju, S., Przytycka, T. M., Allen, A. M., Goodwin, S. F., Berry, C. W., Fuller, M. T., WHITE-Cooper, H., Matunis, E. L., Dinardo, S., Galenza, A., O’brien, L. E., Dow, J. A. T., Jasper, H., Oliver, B., Perrimon, N., Deplancke, B., Quake, S. R., Luo, L., Aerts, S., Agarwal, D., AHMED-Braimah, Y., Arbeitman, M., Ariss, M. M., Augsburger, J., Ayush, K., Baker, C. C., Banisch, T., Birker, K., Bodmer, R., Bolival, B., Brantley, S. E., Brill, J. A., Brown, N. C., Buehner, N. A., Cai, X. T., CARDOSO-Figueiredo, R., Casares, F., Chang, A., Clandinin, T. R., Crasta, S., Desplan, C., Detweiler, A. M., Dhakan, D. B., Donà, E., Engert, S., FLOC’hlay, S., George, N., GONZÁLEZ-Segarra, A. J., Groves, A. K., Gumbin, S., Guo, Y., Harris, D. E., Heifetz, Y., Holtz, S. L., Horns, F., Hudry, B., Hung, R.-J., Jan, Y. N., Jaszczak, J. S., Jefferis, G. S. X. E., Karkanias, J., Karr, T. L., Katheder, N. S., Kezos, J., Kim, A. A., Kim, S. K., Kockel, L., Konstantinides, N., Kornberg, T. B., Krause, H. M., Labott, A. T., Laturney, M., Lehmann, R., Leinwand, S., Li, J., Li, J. S. S., Li, K., et al. 2022. Fly Cell Atlas: A single-nucleus transcriptomic atlas of the adult fruit fly. Science, 375, eabk2432.

Lin, H. H., Kuang, M. C., Hossain, I., Xuan, Y., Beebe, L., Shepherd, A. K., Rolandi, M. & Wang, J. W. 2022. A nutrient-specific gut hormone arbitrates between courtship and feeding. Nature, 602, 632–638.

Liu, X., Nagy, P., Bonfini, A., Houtz, P., Bing, X. L., Yang, X. & Buchon, N. 2022a. Microbes affect gut epithelial cell composition through immune-dependent regulation of intestinal stem cell differentiation. Cell Rep, 38, 110572.

Liu, Y., Li, J. S. S., Rodiger, J., Comjean, A., Attrill, H., Antonazzo, G., Brown, N. H., Hu, Y. & Perrimon, N. 2022b. FlyPhoneDB: an integrated web-based resource for cell-cell communication prediction in Drosophila. Genetics, 220.

Manion, J., Musser, M. A., Kuziel, G. A., Liu, M., Shepherd, A., Wang, S., Lee, P.-G., Zhao, L., Zhang, J., Marreddy, R. K. R., Goldsmith, J. D., Yuan, K., Hurdle, J. G., Gerhard, R., Jin, R., Rakoff-Nahoum, S., Rao, M. & Dong, M. 2023. C. difficile intoxicates neurons and pericytes to drive neurogenic inflammation. Nature, 622, 611–618.

Martin, G. R., Wallace, L. E., Hartmann, B., Holst, J. J., Demchyshyn, L., Toney, K. & Sigalet, D. L. 2005. Nutrient-stimulated GLP-2 release and crypt cell proliferation in experimental short bowel syndrome. Am J Physiol Gastrointest Liver Physiol, 288, G431–8.

Mccarthy, N., Manieri, E., Storm, E. E., Saadatpour, A., Luoma, A. M., Kapoor, V. N., Madha, S., Gaynor, L. T., Cox, C., Keerthivasan, S., Wucherpfennig, K., Yuan, G. C., DE Sauvage, F. J., Turley, S. J. & Shivdasani, R. A. 2020. Distinct Mesenchymal Cell Populations Generate the Essential Intestinal BMP Signaling Gradient. Cell Stem Cell, 26, 391–402.e5.

Medina, A., Bellec, K., Polcowñuk, S. & Cordero, J. B. 2022. Investigating local and systemic intestinal signalling in health and disease with Drosophila. Dis Model Mech, 15.

Mishima, T., Ito, Y., Hosono, K., Tamura, Y., Uchida, Y., Hirata, M., Suzsuki, T., Amano, H., Kato, S., Kurihara, Y., Kurihara, H., Hayashi, I., Watanabe, M. & Majima, M. 2011. Calcitonin gene-related peptide facilitates revascularization during hindlimb ischemia in mice. Am J Physiol Heart Circ Physiol, 300, H431–9.

Morris, O. & Jasper, H. 2021. Reactive Oxygen Species in intestinal stem cell metabolism, fate and function. Free Radic Biol Med, 166, 140–146.

Niec, R. E., Chu, T., Schernthanner, M., Gur-Cohen, S., Hidalgo, L., Pasolli, H. A., Luckett, K. A., Wang, Z., Bhalla, S. R., Cambuli, F., Kataru, R. P., Ganesh, K., Mehrara, B. J., PE’er, D. & Fuchs, E. 2022. Lymphatics act as a signaling hub to regulate intestinal stem cell activity. Cell Stem Cell, 29, 1067–1082.e18.

Ovington, K. S. 1985. Dose-dependent relationships between Nippostrongylus brasiliensis populations and rat food intake. Parasitology, 91 (Pt 1), 157–67.

PALIKUQI B, RISPAL J, Reyes Ea, Vaka D, Boffelli D, Klein O. Lymphangiocrine signals are required for proper intestinal repair after cytotoxic injury. Cell Stem Cell, 29, 1262–1272.e5.

Parikh, K., Antanaviciute, A., Fawkner-Corbett, D., Jagielowicz, M., Aulicino, A., Lagerholm, C., Davis, S., Kinchen, J., Chen, H. H., Alham, N. K., Ashley, N., Johnson, E., Hublitz, P., Bao, L., Lukomska, J., Andev, R. S., Björklund, E., Kessler, B. M., Fischer, R., Goldin, R., Koohy, H. & Simmons, A. 2019. Colonic epithelial cell diversity in health and inflammatory bowel disease. Nature, 567, 49–55.

Parsons, B. & Foley, E. 2013. The Drosophila platelet-derived growth factor and vascular endothelial growth factor-receptor related (Pvr) protein ligands Pvf2 and Pvf3 control hemocyte viability and invasive migration. J Biol Chem, 288, 20173–83.

Perochon, J., Yu, Y., Aughey, G. N., Medina, A. B., Southall, T. D. & Cordero, J. B. 2021. Dynamic adult tracheal plasticity drives stem cell adaptation to changes in intestinal homeostasis in Drosophila. Nat Cell Biol, 23, 485–496.

Petsakou, A., Liu, Y., Liu, Y., Comjean, A., Hu, Y. & Perrimon, N. 2023. Epithelial Ca(2+) waves triggered by enteric neurons heal the gut. bioRxiv.

Russell, F. A., King, R., Smillie, S. J., Kodji, X. & Brain, S. D. 2014. Calcitonin gene-related peptide: physiology and pathophysiology. Physiol Rev, 94, 1099–142.

Sasaki, M., Fitzgerald, A. J., Mandir, N., Sasaki, K., Wright, N. A. & Goodlad, R. A. 2001. Glicentin, an active enteroglucagon, has a significant trophic role on the small intestine but not on the colon in the rat. Aliment Pharmacol Ther, 15, 1681–6.

Sciola, V., Massironi, S., Conte, D., Caprioli, F., Ferrero, S., Ciafardini, C., Peracchi, M., Bardella, M. T. & Piodi, L. 2009. Plasma chromogranin a in patients with inflammatory bowel disease. Inflamm Bowel Dis, 15, 867–71.

Scopelliti, A., Bauer, C., Yu, Y., Zhang, T., Kruspig, B., Murphy, D. J., Vidal, M., Maddocks, O. D. K. & Cordero, J. B. 2018. A Neuronal Relay Mediates a Nutrient Responsive Gut/Fat Body Axis Regulating Energy Homeostasis in Adult Drosophila. Cell Metab.

Scopelliti, A., Bauer, C., Yu, Y., Zhang, T., Kruspig, B., Murphy, D. J., Vidal, M., Maddocks, O. D. K. & Cordero, J. B. 2019. A Neuronal Relay Mediates a Nutrient Responsive Gut/Fat Body Axis Regulating Energy Homeostasis in Adult Drosophila. Cell Metab, 29, 269–284.e10.

Scopelliti, A., Cordero, J. B., Diao, F., Strathdee, K., White, B. H., Sansom, O. J. & Vidal, M. 2014. Local control of intestinal stem cell homeostasis by enteroendocrine cells in the adult Drosophila midgut. Curr Biol, 24, 1199–211.

Stzepourginski, I., Nigro, G., Jacob, J.-M., Dulauroy, S., Sansonetti, P. J., Eberl, G. & Peduto, L. 2017. CD34^+^ mesenchymal cells are a major component of the intestinal stem cells niche at homeostasis and after injury. Proceedings of the National Academy of Sciences, 114, E506–E513.

Tamamouna, V., Rahman, M. M., Petersson, M., Charalambous, I., Kux, K., Mainor, H., Bolender, V., Isbilir, B., Edgar, B. A. & Pitsouli, C. 2021. Remodelling of oxygen-transporting tracheoles drives intestinal regeneration and tumorigenesis in Drosophila. Nat Cell Biol, 23, 497–510.

Tauc, H. M., Rodriguez-Fernandez, I. A., Hackney, J. A., Pawlak, M., Ronnen Oron, T., Korzelius, J., Moussa, H. F., Chaudhuri, S., Modrusan, Z., Edgar, B. A. & Jasper, H. 2021. Age-related changes in polycomb gene regulation disrupt lineage fidelity in intestinal stem cells. Elife, 10.

Toda, M., Suzuki, T., Hosono, K., Kurihara, Y., Kurihara, H., Hayashi, I., Kitasato, H., Hoka, S. & Majima, M. 2008. Roles of calcitonin gene-related peptide in facilitation of wound healing and angiogenesis. Biomed Pharmacother, 62, 352–9.

Van Marle, G., Sharkey, K. A., Gill, M. J. & Church, D. L. 2013. Gastrointestinal viral load and enteroendocrine cell number are associated with altered survival in HIV-1 infected individuals. PLoS One, 8, e75967.

Veenstra, J. A., Agricola, H. J. & Sellami, A. 2008. Regulatory peptides in fruit fly midgut. Cell Tissue Res, 334, 499–516.

Worthington, J. J., Reimann, F. & Gribble, F. M. 2018. Enteroendocrine cells-sensory sentinels of the intestinal environment and orchestrators of mucosal immunity. Mucosal Immunol, 11, 3–20.

Xu, H., Ding, J., Porter, C. B. M., Wallrapp, A., Tabaka, M., Ma, S., Fu, S., Guo, X., Riesenfeld, S. J., Su, C., Dionne, D., Nguyen, L. T., Lefkovith, A., Ashenberg, O., Burkett, P. R., Shi,

H. N., Rozenblatt-Rosen, O., Graham, D. B., Kuchroo, V. K., Regev, A. & Xavier, R. J. 2019. Transcriptional Atlas of Intestinal Immune Cells Reveals that Neuropeptide α-CGRP Modulates Group 2 Innate Lymphoid Cell Responses. Immunity, 51, 696–708.e9.

Yang, S., Gaafar, S. M. & Bottoms, G. D. 1990. Effects of multiple dose infections with Ascaris suum on blood gastrointestinal hormone levels in pigs. Vet Parasitol, 37, 31–44.

Zhang, X., Zhang, H., Shen, B. & Sun, X. F. 2019. Chromogranin-A Expression as a Novel Biomarker for Early Diagnosis of Colon Cancer Patients. Int J Mol Sci, 20.

Zhou, J. & Boutros, M. 2023. Intestinal stem cells and their niches in homeostasis and disease. Cells & Development, 175, 203862.

Zissimopoulos, A., Vradelis, S., Konialis, M., Chadolias, D., Bampali, A., Constantinidis, T., Efremidou, E. & Kouklakis, G. 2014. Chromogranin A as a biomarker of disease activity and biologic therapy in inflammatory bowel disease: a prospective observational study. Scand J Gastroenterol, 49, 942–9.

