## Supplementary figures and images for "Neuroendocrine Control of Intestinal Regeneration Through the Vascular Niche in *Drosophila*"

### Supplementary Figure 1-5

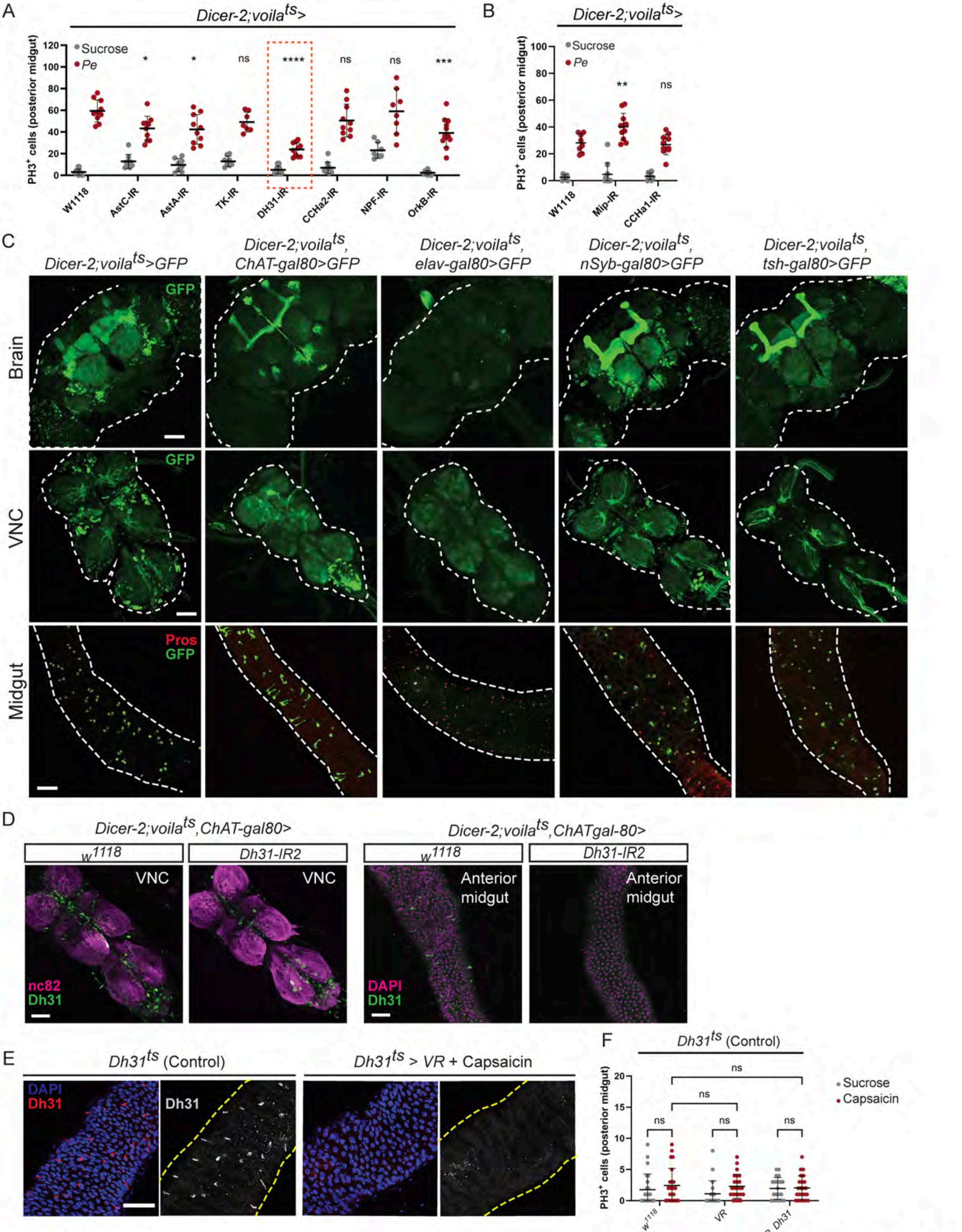


Medina *et al.* Figure S1

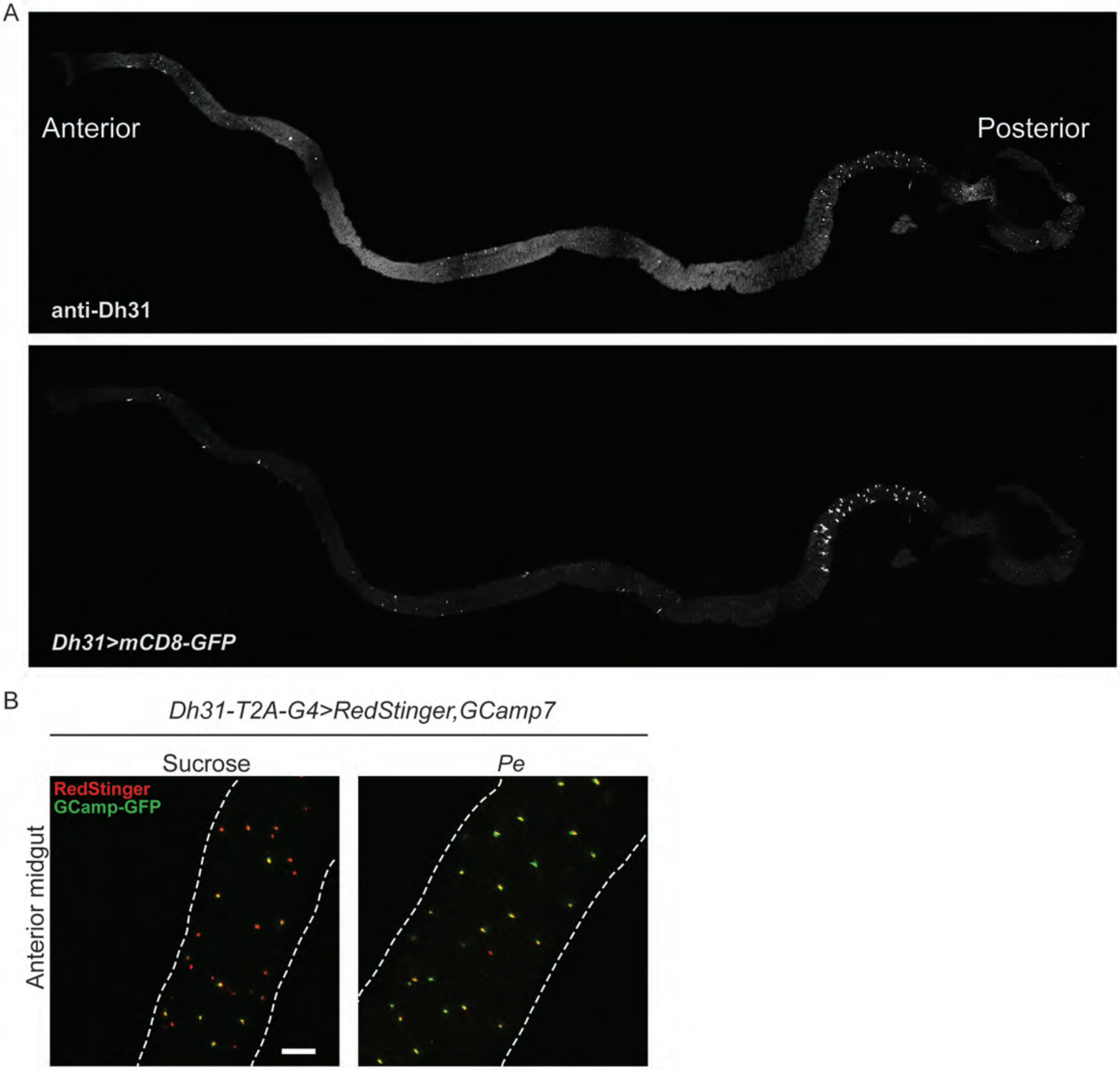


Medina *et al.* Figure S2

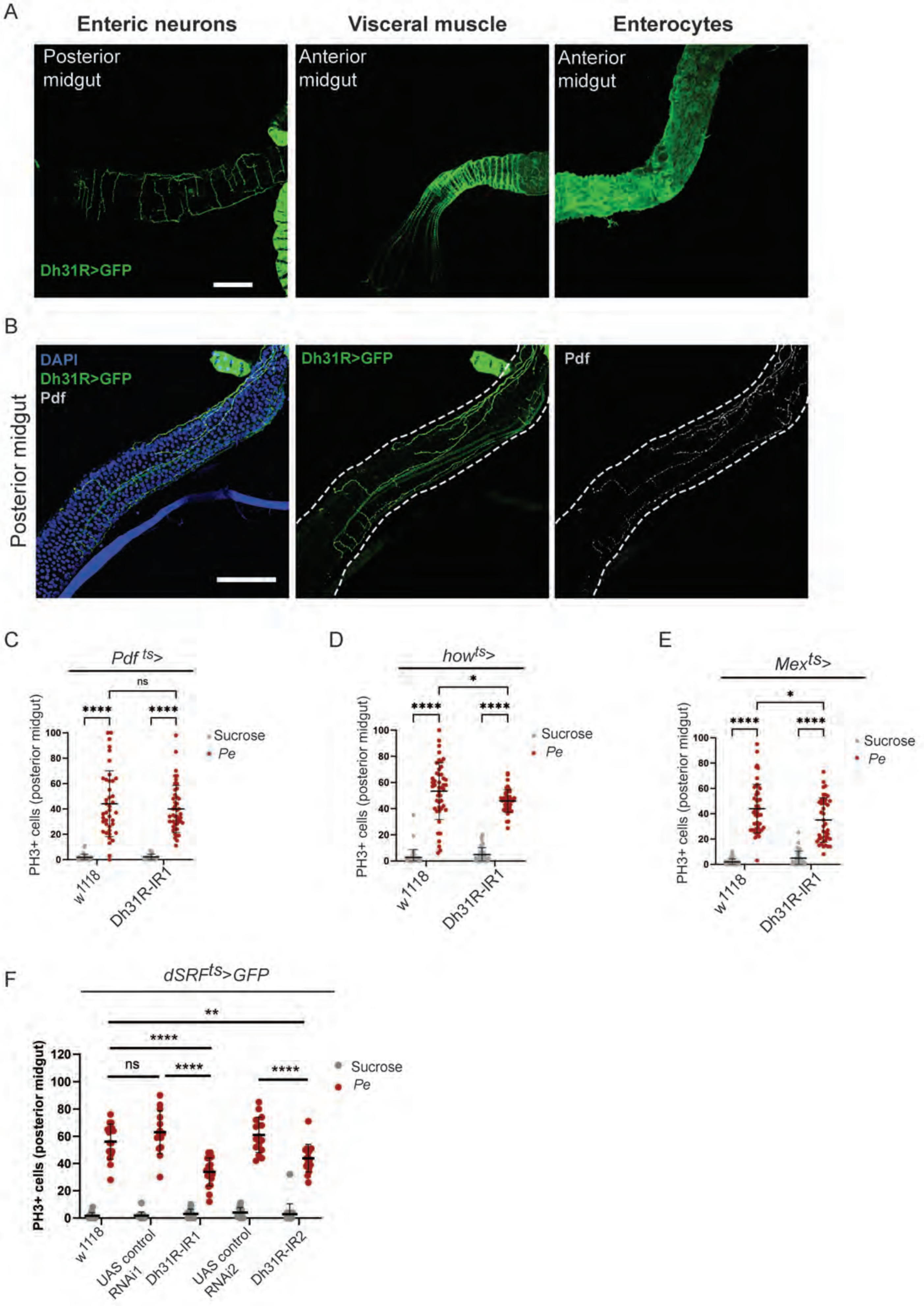


Medina *et al.* Figure S3

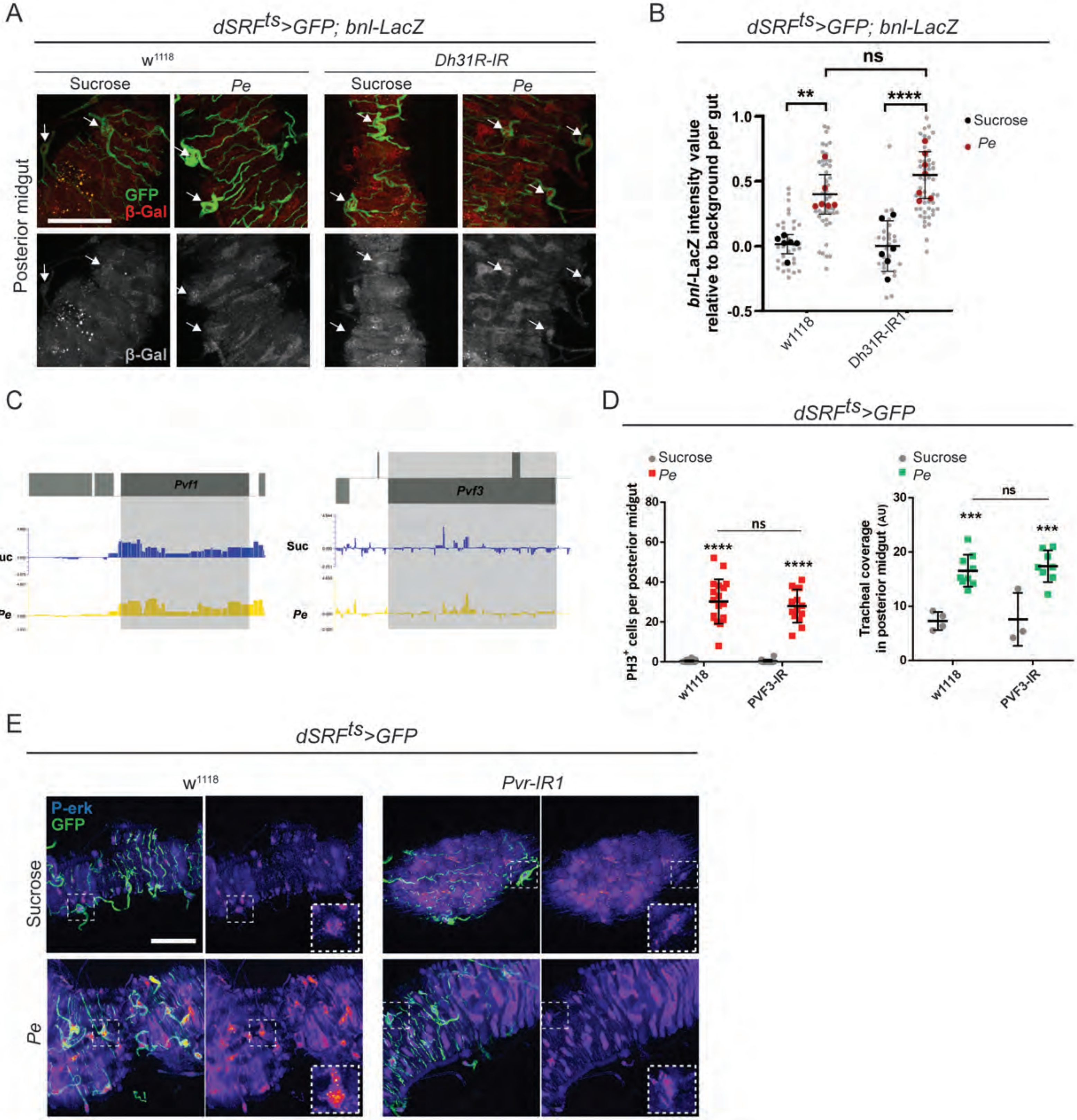


Medina *et al.* Figure S4

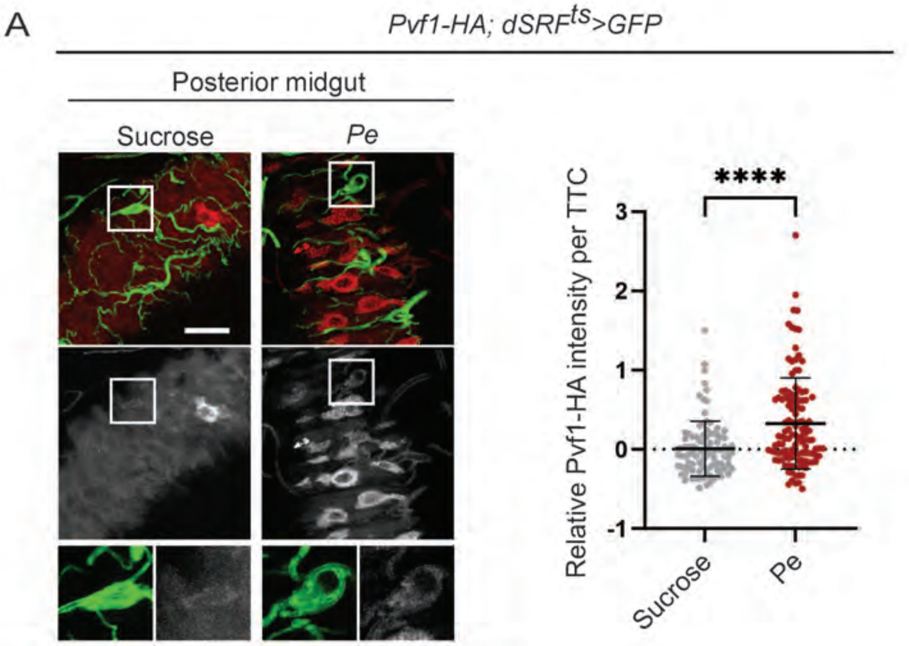
